# Element partitioning, element localisation and transcriptome responses of date palms exposed to NaCl

**DOI:** 10.1101/2024.12.07.627310

**Authors:** Paula Pongrac, Sina Fischer, Gladys Wright, Jacqueline A. Thompson, Mitja Kelemen, Konrad Neugebauer, Lawrie K. Brown, Katarina Vogel-Mikuš, Martin Šala, Hamed A. El-Serehy, Philip J. White

## Abstract

Date palm (*Phoenix dactylifera* L.) is an economically important fruit crop in many (semi)arid regions. Large intra-species variation in susceptibility to NaCl has been reported. Controlled experiments were conducted to evaluate sodium (Na) and chlorine (Cl) partitioning in two-year old plants of six date palm varieties and to determine tissue location of Na and Cl and transcriptome responses to NaCl of two varieties. The largest Na concentrations were found in secondary roots, especially under NaCl treatment. By contrast, only small differences in within-plant partitioning of Cl were observed with less differences between varieties than for Na. Variety Sultana had the largest Na translocation factor. The variety Khalas exhibited better restriction of Na mobility, which may be a consequence of Na hotspots in roots, presumably immobilising Na in roots, which were absent in Sultana. In the root, shoot base and lower part of leaves, Cl accumulated in fibre bands and sclerenchyma sheaths of vascular bundles. RNAseq analysis revealed that Sultana and Khalas respond very differently to NaCl: Khalas modified expression of NaCl-specific genes in both roots and leaves, while Sultana responded manly in leaves. Gene ontology analysis showed that the expression changes for genes linked with photosynthesis and carbohydrate metabolism predominated as a response to NaCl. In Khalas, differentially expressed genes encoding for metal transporters and those reported to be involved in heat, salt and osmotic stress response were found. Large variability in Na and Cl accumulation in different date palm varieties may have practical agronomical consequences.

## INTRODUCTION

Date palm (*Phoenix dactylifera*) is an economically important fruit crop in many arid and semiarid regions of the world (Chao and Krueger 2007; Al-Khayri et al., 2015a, b; Yaish and Kumar 2015). Although it is classified as a salt-tolerant crop (Maas and Grattan 1999; Ramoliya and Pandey 2003; Alrasbi et al., 2010; Alhammadi and Kurup 2012; Yaish and Kumar 2015; Hazzouri et al., 2020), there is considerable variation among date palm varieties in their susceptibility to NaCl (Alhammadi and Edward 2009; Alrasbi et al., 2010; Al Kharusi et al., 2017; Al-Muaini et al., 2019; Al-Qurainy et al., 2020; Al-Dakheel et al., 2022). This can result in large economic losses when susceptible varieties are cultivated on saline soils (Tripler et al., 2011; Alhammadi and Kurup 2012; Yaish and Kumar 2015; Al-Muaini et al., 2019; Hazzouri et al., 2020).

Increasing salinity in the rhizosphere decreases stomatal conductance, transpiration and CO_2_ assimilation in date palm (Ramoliya and Pandey 2003; Sperling et al., 2014; Yaish and Kumar 2015; Al Kharusi et al., 2017; Al-Harrasi et al., 2018; Al-Muaini et al., 2019; Jana et al., 2019; Du et al., 2021). However, a low stomatal conductance can be maintained in most varieties even when grown in saline environments (Ramoliya and Pandey 2003; Yaish et al., 2017; Al Kharusi et al., 2017; Al-Muaini et al., 2019; Al Kharusi et al., 2019a; Jana et al., 2019; Du et al., 2021), although photosynthesis and growth are greatly curtailed in salt-sensitive varieties (Alhammadi and Edward 2009; Al Kharusi et al., 2017). Date palm varieties have been organised in four groups depending on their yield at high salinity, but inconsistently. For example, variety Khalas has been considered to be a relativley salt tolerant variety (Yaish and Kumar 2015), less tolerant (Alrasbi et al., 2010; Al Kharusi et al., 2017) and salt intolerant (Al-Muaini et al., 2019) depending on the cultivars being compared.

Salinity tolerance in plants is largely based upon (a) sodium (Na) exclusion by roots, (b) restricting Na accumulation in leaves, (c) biochemical compensation for osmotic stress and (d) “metabolic tolerance” (Munns and Tester. 2008). Although the mechanistic basis for salinity tolerance in date palm has been investigated in several recent studies using ‘omics’ approaches (Radwan et al., 2015; El Rabey et al., 2016; Al Kharusi et al., 2017, 2019 *a, b*; Patankar et al., 2018; Jana et al., 2019; Al-Harrasi et al., 2020; Du et al., 2021; Mueller et al., 2023) and functional genomics analysis approach (Yaish et al., 2017; Patankar et al., 2019; Jana et al., 2021; Jana and Yaish 2020; Rahman et al., 2022) information on the responses of date palms to salinity is still limited (Yaish and Kumar 2015; Hazzouri et al., 2020).

It is thought that differences in salinity tolerance among date palm varieties are based primarily on their contrasting abilities to limit Na uptake and accumulation in leaves, whilst maintaining sufficient tissue potassium (K) concentrations (Alhammadi and Edward 2009; Alrasbi et al., 2010; Al Kharusi et al., 2017, 2019 *a, b*). Maintaining a large cytosolic K/Na quotient is one of the key features of salt tolerance (Munns and Tester 2008; Rubio et al., 2020). In addition, the accumulation of compatible solutes to offset osmotic stress (Du et al., 2021), and the mitigation of oxidative stress caused by increased formation of reactive oxygen species due to the reduced rate of photosynthesis (Munns and Tester 2008), also contribute to salinity tolerance in date palm (Al Kharusi et al., 2019b). Recently Mueller et al., (2023) have provided ionomic, transcriptomic, proteomic and metabolomic evidence on the salt tolerance of date palm variety Khalas. They showed that tolerance in Khalas is based on abilities (1) to avoid ionic stresses, (2) to avoid osmotic stress and (3) to mitigate oxidative damage (Mueller et al., 2023). The following mechanisms behind these abilities were proposed: (1) upregulation of the expression of genes encoding transport proteins that facilitate the exclusion of Na and Cl from roots and restrict their translocation to leaves, (2) accumulation of compatible solutes, primarily amino acids, in both roots and shoots and (3) by scavenging reactive oxygen species, respectively (Mueller et al., 2023).

In this study, element partitioning in two-year old plants of six different date palm varieties exposed to NaCl is reported. Further, in two date palm varieties with contrasting abilities to restrict Na translocation, tissue location of Na and Cl (and other essential elements) is shown in detail, which is further expanded by comparative transcriptome responses to NaCl in the two contrasting varieties. The following hypothesese were tested: 1) there is significant difference in element partitioning in selected date palm varieties, 2) differences in Na translocation can be explained by different Na allocation mechanism(s), and 3) differences in Na translocation can be explained by different expression pattern of NaCl-responsive genes.

## MATERIALS AND METHODS

### Experiment 1. Identification of Varieties with Contrasting Sodium Partitioning Between Roots and Leaves

Twelve two-year old plants of six varieties (Barhee, Khalas, Lulu, Nabut Saif, Sukkary and Sultana) of micro-propagated date palms were obtained from Date Palm Developments Ltd. (Somerset, U.K.). Six plants of each variety were harvested immediately after purchasing (they are referred to as pre-experiment plants). These plants were separated into leaves and roots and processed as described below.

The remaining six plants were first transplanted to 4 L pots filled with standard potting compost. The standard potting compost was made in bulk and comprised 75% peat (Sinclairs Professional Peat, Sinclair Pro, Ellesmere Port, UK) and 25% sand mixed with 0.225 g L^-1^ single superphosphate, 0.4 g L^-1^ ammonium nitrate, 0.75 g L^-1^ potassium nitrate, 2.25 g L^-1^ ground limestone, 2.25 g L^-1^ Magnesian limestone and 0.51 g L^-1^ of a trace element mixture containing 0.16% boron, 0.79% copper, 11.82% iron, 1.97% manganese, and 0.04% molybdenum by weight. Plants were then grown in a glasshouse at the James Hutton Institute, Dundee (U.K.; latitude 56°270 26″N, longitude 3°40 17″W). The glasshouse was set to maintain 25 °C day and 18 °C night temperatures with a day length of 16 h using automatic venting, supplementary heating and additional lighting as described previously (White et al., 2017). The six varieties were grown in a randomised block design with six replicates. Plants were acclimated for two weeks before treatments were applied, namely NaCl or control treatments. Each plant subject to the NaCl treatment was irrigated once with 400 mL of 150 mM NaCl at the beginning of the experiment, then after two, four and six weeks with 400 mL of 300 mM NaCl. Plants that were not subject to the NaCl treatment were irrigated with 400 mL double-distilled water on each occasion. At harvest, seedings were separated into primary root, secondary root, shoot base and leaves.

Roots of all plants were carefully washed in tap water to remove soil debris. All tissues were dried in an oven at 70 °C to a constant weight and their dry weight (DW) was determined. The concentrations of elements in dried plant material were determined by inductively coupled plasma mass spectrometry (ICP-MS) as described below (Elemental Analyses).

### Experiment 2. Tissue-specific Element Localisation Analysis in Khalas and Sultana Varieties

Two-year old plants of micro-propagated date palm varieties (Khalas and Sultana) were obtained from Date Palm Developments Ltd. (Somerset, U.K.). Upon arrival at University of Ljubljana, Biotechnical Faculty, Ljubljana, Slovenia, plants were transplanted to 1.5 L pots filled with a mixture of Compo Sana® Potting Soil for Green Plants and Palms (Compo GmbH, Münster, Germany) and sand (5:2 v/v) and watered to water-holding capacity with double-distilled water. Plants were acclimatised to growth in a controlled environment chamber with 60% humidity, a temperature regime of 25 °C day and 20 °C night, a day length of 16 h, and 250 µmol s^-1^ m^-2^ light intensity at plant height and were then watered as needed. After one month of acclimation, two plants per variety were watered with 100 mL of either double-distilled water (control treatment) or three plants per variety were watered with 150 mM NaCl three times per week for two weeks, followed by 300 mM NaCl two times per week for four weeks (NaCl treatment). Root tips, bases of the roots, bases of the shoot, base of the leaf and tip of the leaf were harvested to map the spatial distributions of Na, chlorine (Cl), K, phosphorus (P), sulphur (S), silicon (Si) and calcium (Ca) within plant tissues as described below (Localisation of Elements) and the concentrations of elements in dried plant material (the separated tissues and the remaining of the roots, base of the shoot and leaves) were determined by ICP-MS, as described below (Elemental Analyses).

### Experiment 3. Effect of NaCl on the Transcriptomes of Roots and Shoots of Khalas and Sultana Varieties

Two-year old plants of micro-propagated date palm varieties (Khalas and Sultana) were obtained from Date Palm Developments Ltd. (Somerset, U.K.) and grown as described for Experiment 2, except that there were five control plants and five NaCl-treated plants per variety. Total RNA was extracted from roots and shoots and RNAseq was performed as described below (Transcriptome Analyses).

### Elemental Analyses

The concentration of elements in dried plant material from Experiments 1 and 2 were determined by ICP-MS following acid digestion. Dried samples were milled to a fine powder using a ceramic ball mill (Retsch MM 200 or Retsch MM 301; Retsch, Haan, Germany) and accurately weighed powdered subsamples (∼50 mg DW) were digested in closed vessels using a microwave digester (MARS Xpress, CEM Microwave Technology, Buckingham, UK) as described by (White et al., 2012). Samples were first digested with 3 mL concentrated HNO_3_ before the addition of 1 mL of 30% H_2_O_2_ to complete digestion. Digested samples were diluted to 50 mL with sterile double-distilled water before element analyses. Total K, Ca, magnesium (Mg), P, S, Na, Cl, iron (Fe), manganese (Mn), zinc (Zn), copper (Cu) and nickel (Ni) concentrations of digested samples were determined by ICP-MS (ELAN DRCe, PerkinElmer, Waltham, MA, USA. [Experiment 1] or Nexion 1000, PerkinElmer, Waltham, MA, USA [Experiment 2]). Blank digestions were performed to determine background concentrations of elements and a tomato leaf standard (Reference 1573a; National Institute of Standards and Technology, NIST, Gaithersburg, MD, USA) was used as an analytical control.

### Localisation of Elements

The spatial distributions of Na, Cl, K, P, S, Si and Ca were determined in cross sections of root tips, base of the root, shoot base, base of the leaves and leaf tips. At harvest, whole plants were removed from the pot and all soil particles were removed before the roots were washed in tap water and with double-distilled water. Plant tissues were excised such that they could fit into the 2 mm opening of a stainless-steel needle. Cryo-fixation, cryo-sectioning to 50 µm thickness, and freeze drying was performed as described previously (Vogel-Mikuš et al., 2014). Localisation of elements within 50 µm cross sections was assessed using micro-particle induced X-ray emission (micro-PIXE) as described previously (Pongrac et al., 2013) at the nuclear microprobe in the Tandetron Accelerator Laboratory of the Jožef Stefan Institute, Ljubljana, Slovenia (Simičič et al., 2002). The sample area ranged from 400 × 400 to 1000 × 1000 µm^2^ at 2-4 µm lateral resolution. Emitted X-rays were detected by a silicon drift detector (RaySpec Ltd, High Wycombe, Buckinghamshire, UK). The collected spectra were analysed and tissue-specific elemental maps were generated in GeoPIXE II (Ryan, 2000). A Gaussian function at 1.5 was applied to smooth the images. Co-localisation maps were generated in Fiji software (Schindelin et al., 2012) with the Color>Merge Channels function on the numerical concentration matrices exported as comma-separated values files from GeoPIXE II. Associations between Na and K and between Na and Cl were captured from the representative distribution maps in root tip, base of the root, base of the leaf and tip of the leaf using Element Association feature in GeoPIXEII.

### Transcriptome Analyses

Roots and shoots were frozen in liquid nitrogen and kept at -80 °C until RNA extraction. Plant material was first homogenised by grinding manually in liquid nitrogen, then a subsample was taken and further pulverised before extraction. The RNAqueousTM kit for total RNA isolation (Thermo Fisher Scientific, MA, USA) was used according to the manufacturer’s instructions. RNA integrity and concentration were confirmed using a 1% (w/v) agarose gel. RNA was sent to BGI Hong Kong (https://www.bgi.com/global) for sequencing. Single-end sequencing was performed on a DNBseq platform. Library preparation, sequencing, alignment and bioinformatics were performed by BGI according to their standard procedures. Each sample generated an average of 4.33 Gb of sequence and had an average mapping ratio of 84.04% with the reference genome. Genes were mapped with 100% accuracy and 28,027 genes were identified of which 2,355 were novel genes and 17,570 were novel transcripts. After filtering the raw reads using SOAPnuke (version1.5.2), which removed adaptors and unknown bases as well as low-quality reads, clean reads were mapped to the reference genome (NCBI_GCF_009389715.1_palm_55x_up_171113_PBpolish2nd_filt_p) using HISAT2 (version 2.0.4). StringTie (version 1.0.4) (Pertea et al., 2015) was used to predict transcripts and Cuffcompare (Trapnell et al., 2012) to identify new transcripts. Finally, CPC (version 0.9- r2) (Kong et al., 2007) was used to predict the coding potential of the new transcripts and the subsequently selected new transcripts were merged with the reference transcripts. Gene expression was calculated using RSME (version 1.2.5) (Li and Dewey, 2011). Differentially expressed genes were extrapolated by using DEseq2. The expression was given as Transcripts Per Kilobase Million (TPM). A fold change was determined from the average expression in two different groups and DEGs were then filtered for a fold change ≥2 and an adjusted p value ≤0.05.

### Statistical Analysis

Data on element concentrations were expressed as means ± standard error of the mean from n observations. One-Way Analysis of Variance (ANOVA) was used on raw data and when significant, Holm-Sidak post-hoc test at p<0.05 was used to determine statistical differences between means. For pairwise comparisons, Student t-test was performed with the level of significance indicated in the text. Graphs were generated and statistics for element composition were performed in SigmaPlot v 12.0 (Systat Software, San Jose, CA, USA).

## RESULTS AND DISCUSSION

### Palm Varieties with Different NaCl Sensitivity Show Different Element Partitioning

Increasing NaCl in irrigation water increased both Na and Cl concentrations in the roots and the leaves of six date palm varieties studied (**Figure 1**). Concentration of Cl was much larger than Na in all date palm parts at both treatments. However, the relative increase in Na concentration (exceeding 3-fold increase in all organs) was greater than that of Cl (remaining below 1.5 in all organs) which is consistent with previous studies (e.g. Furr and Armstrong 1962; Alrasbi et al., 2010; Du et al., 2021; Mueller et al., 2023) and suggests that roots of date palm could restrict Cl entry to the plant better than Na in saline environments. Despite the observed increase in Na and Cl concentrations, there was no biomass penalty with increased NaCl in any of the date palm varieties (**Figure S1A**), indicating an absence of Na or Cl toxicity in the experiment. There was, however, a considerable difference in biomass increase during the experiment for different varieties with the largest biomass accumulation observed for varieties Lulu, Nabut Saif and Sukkary (**Figure S1B**). Variety Lulu has been reported to have the highest fruit yield, even at increased salinity, among 18 date palm varieties investigated (Al-Dakheel et al., 2022). However, high-yielding and salt tolerant was also variety Barhi, while Sukkari was grouped into high-yielding varieties with sensitivity to increased salinity, and variety Nabtat Saif having low yield potential with low salinity tolerance in a long-term experiment (Al-Dakheel et al., 2022).

**Figure 1.**
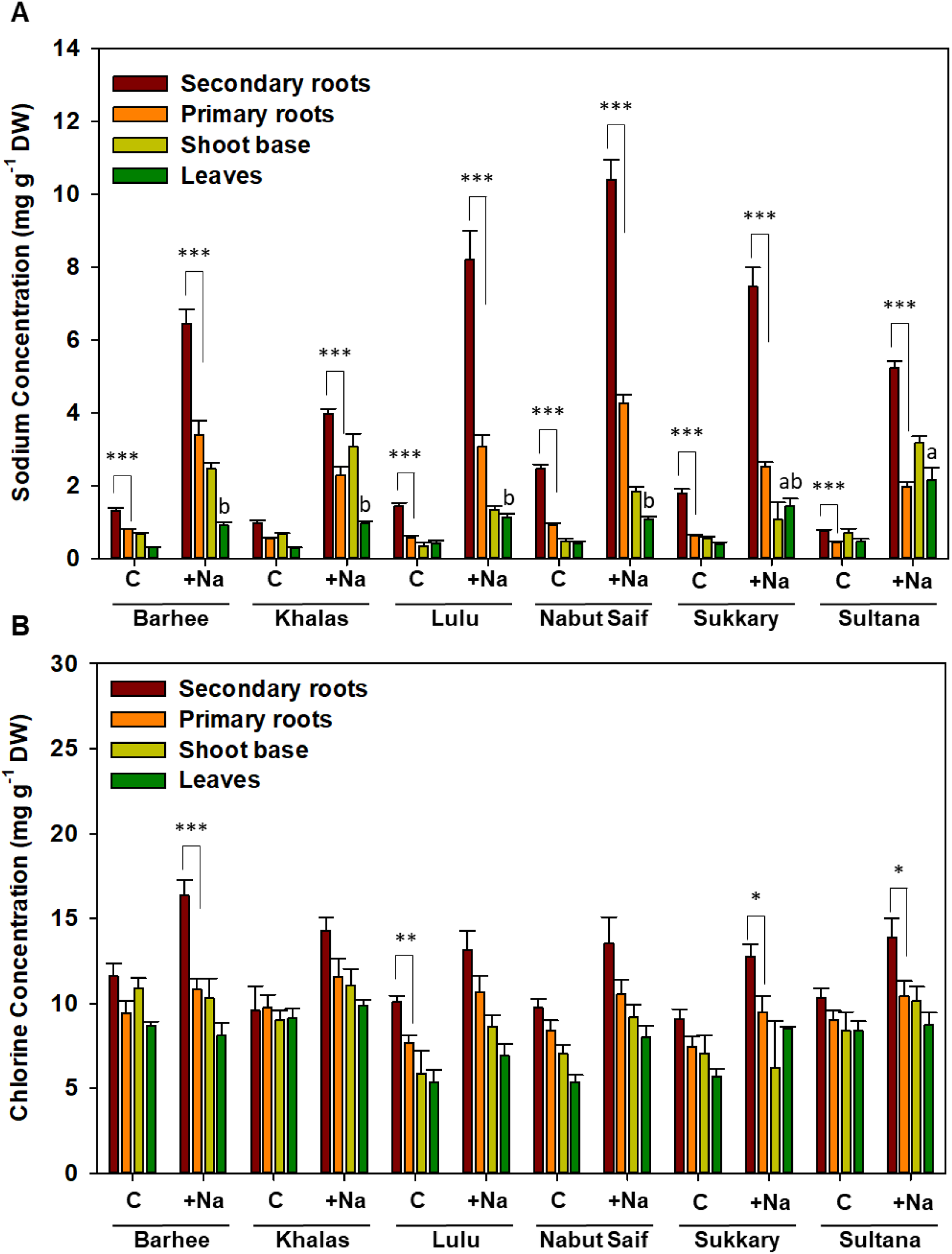
Sodium (A) and chlorine (B) concentration in different parts of six date palm (*Phoenix dactylifera*) varieties. Plants were irrigated with double-distilled water (Control, C) or once with 400 mL of 150 mM NaCl at the beginning of the experiment, then after two, four and six weeks with 400 mL of 300 mM NaCl (+Na); Experiment 1. Shown are averages ± standard errors (n=2-6). Asterisks indicate significant differences as determined by Student t-test at p<0.05 (*), p<0.01 (**) and p<0.001 (***). Different letters above columns indicate significant differences between means for different varieties under salinity (One-Way ANOVA and Holm-Sidak post-hoc at p<0.05); DW, dry weight.

In all varieties the Na concentrations in secondary roots were larger than those in primary roots (t-test at p<0.05). This was seen in both treatments (**Figure 1A**). It is consistent with recent reports on larger Na concentrations in root tips than other parts of the root in date palm Khalas (Mueller et al., 2023). Thus, a bulk of numerous secondary roots, each with a root tip, will accumulate more Na than a primary root with a single root tip. It is likely that the absence of a Casparian strip at the root tip, which controls selective ion transport to the stele, is responsible for this phenotype (Peterson and Lefcourt 1990; Baxter et al., 2009). Significant differences in Na concentrations in secondary roots between varieties indicate that they differ in their ability to exclude, export or translocate Na from their roots. Varieties Khalas and Sultana were best able to restrict Na uptake by roots, judged by their smallest Na concentrations in secondary roots at NaCl treatment (**Figure 1A**). The ability to restrict Na concentrations in plant roots exposed to NaCl has been considered a key trait in NaCl tolerance (Munns and Tester 2008). By contrast to Na concentrations, the Cl concentrations in secondary roots of date palms were not larger than those in primary roots with few exceptions (**Figure 1B**).

There were smaller concentrations of Na and Cl in roots and leaves of pre-experiment plants than in roots (average across the primary and secondary roots; **Table S1**) and leaves of plants from Experiment 1 (**Figure S2**), which indicates increase in Na and Cl concentrations with plant growth and NaCl treatment. Before the start of Experiment 1, there was no differences in Na and Cl concentrations between varieties, which suggest that differences in Na and Cl concentration observed in Experiment 1 were due to genetic differences among varieties, which supports our first hypothesis that there is significant difference in Na and Cl partitioning in selected date palm varieties.

Salt tolerant plants are believed to exhibit small translocation factors for Na (a quotient between Na concentration in the leaves over roots) when exposed to salinity (Munns and Tester 2008; Al Kharusi et al., 2017, 2019a). In pre-experiment plants the Na translocation factor was large and it decreased during the Experiment 1, but in variety Sultana this decrease was smaller than in other varieites (**Figure 2A**). Regression analysis between root and leaf Na concentration also revealed a different behaviour of Sultana. This variety had larger Na concentration in leaves at relatively small root Na concentration when grown in saline conditions compared to other varieties (**Figure 2B**). Due to its poor restriction of Na mobility within this variety, Sultana can be considered a salt-intolerant variety in line with previous observations (Al Kharusi et al., 2017, 2019b).

**Figure 2.**
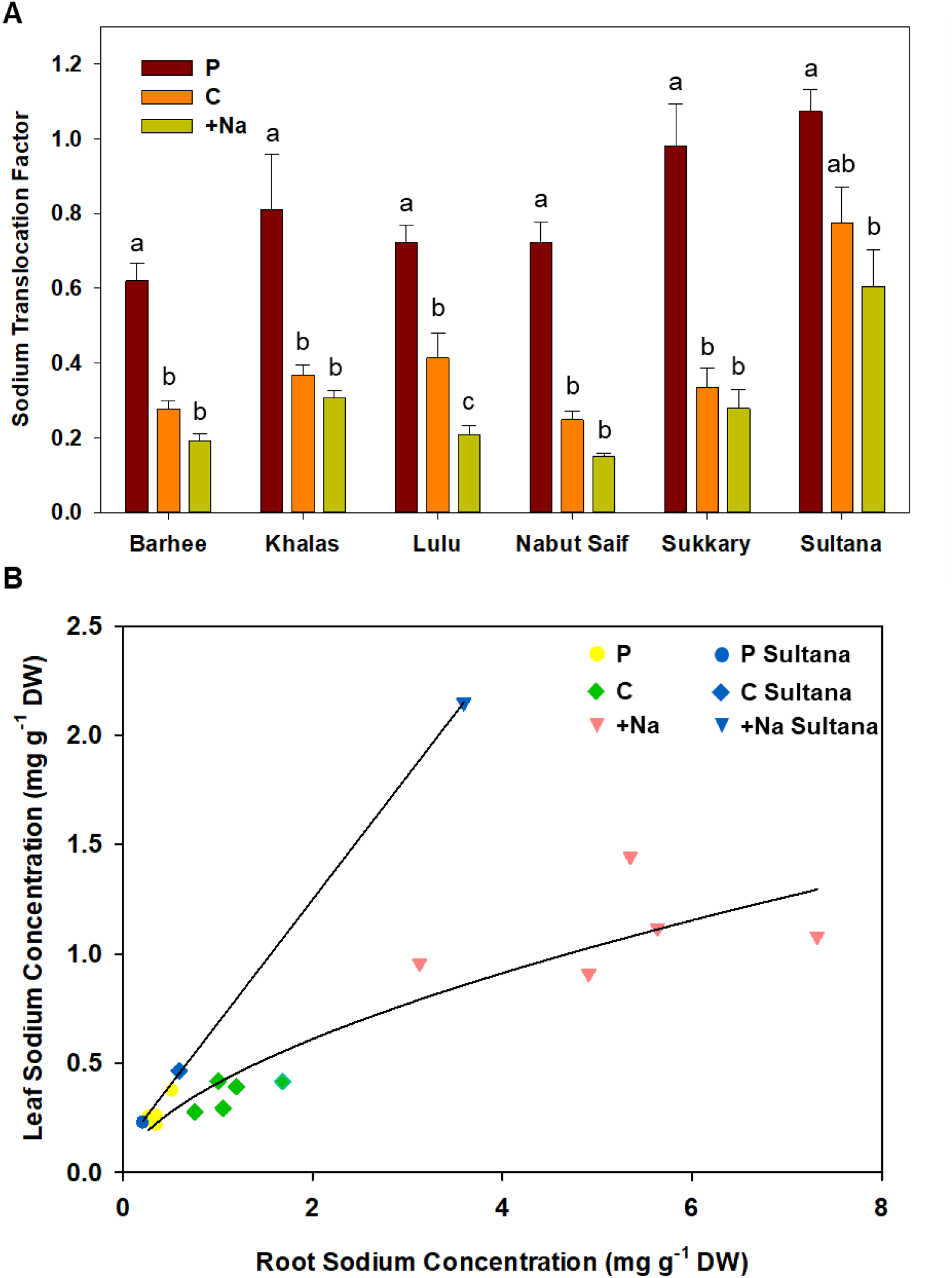
Sodium translocation factors (leaf/ root sodium concentration quotient; A) and regression curves between root and leaf sodium concentration (B) in six date palm (*Phoenix dactylifera*) varieties. Plants were analysed immediately after purchasing (pre-experiment plants; P), irrigated with double-distilled water (Control, C) or once with 400 mL of 150 mM NaCl at the beginning of the experiment, then after two, four and six weeks with 400 mL of 300 mM NaCl (+Na); Experiment 1. Shown are averages ± standard errors (A) and averages (B) of n=2-6. Different letters above columns indicate significant differences between means for each variety separately (One Way ANOVA test and Holm-Sidak post-hoc test at p <0.05); DW, dry weight.

The restricted accumulation of Na in leaves should, in tolerant plants (Munns and Tester 2008; Al Kharusi et al., 2017, 2019b), be accompanied by maintenance of sufficient leaf K concentration. This could potentially result in a larger K/Na quotient than in salt-sensitive varieties when grown in saline environments. Varieties Sultana, Barhee and Khalas had the largest leaf K concentrations (in both non-saline and saline conditions) compared to other varieties (**Figure 3A**). Larger leaf K concentration for variety Barhee is consistent with previous reports (Perveen et al., 2013). Curiously, other studies have observed smaller shoot K concentrations in Barhee and Khalas than in other varieties when grown in non-saline conditions, but a large increase in their shoot K concentrations in response to saline conditions (Alrasbi et al., 2010; Al Kharusi et al., 2017). All date palm varieties in our study maintained their leaf K concentrations when grown in saline conditions compared to non-saline concentrations (**Figure 3A**). This was observed previously for many palm varieties including Barhee, Fard, Khalas, Zabad and Umsila (Al Kharusi et al., 2017, 2019a; Jana et al., 2019; Mueller et al., 2023). One consequence of differences in Na and K concentrations among varieties, and their contrasting responses to NaCl, was a change in the K/Na quotient (**Figure 3B**). The K/Na quotient decreased with increasing NaCl in all plant parts and in all date palm varieties because Na concentrations were increased overall (**Figure 3B**). This was in line with previous reports (Alhammadi and Edward 2009). Similarly, the K/Na quotient tended to be larger in leaves than in roots, particularly secondary roots, in all varieties as reported by Alhammadi and Edward (2009). Secondary roots had smaller K/Na quotient than primary root, except in Barhee and Sukkary under NaCl treatment (**Figure 3B**), due to very large Na concentrations in secondary roots.

**Figure 3.**
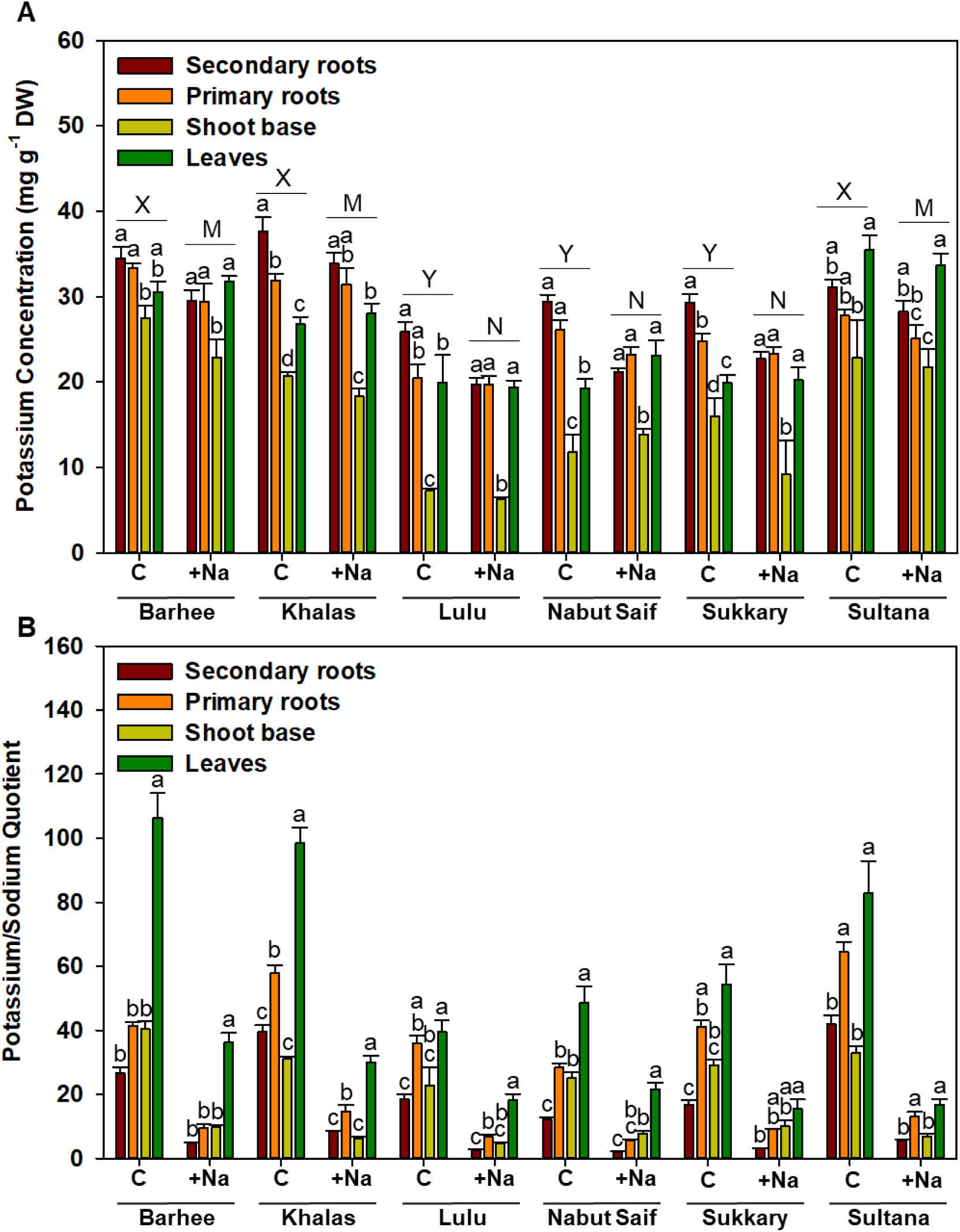
Potassium (A) concentration and potassium /sodium quotient (B) in different parts of six date palm (*Phoenix dactylifera*) varieties. Plants were irrigated with double-distilled water (Control, C) or once with 400 mL of 150 mM NaCl at the beginning of the experiment, then after two, four and six weeks with 400 mL of 300 mM NaCl (+Na); Experiment 1. Shown are averages ± standard errors (n=2-6). Different lowercase letters above columns indicate significant differences between means for different varieties for each treatment separately and upper case letters indicate significant differences between varieties for control treatment (X, Y) and saline treatement (M, N( separately (One-Way ANOVA and Holm-Sidak post-hoc at p<0.05); DW, dry weight.

There was no apparent effect of NaCl on the concentration of any essential element measured in either roots or leaves of the palm varieties, but there were substantial differences among date palm varieties (**Figures S3-6**) in line with previous reports (Kolsi-Benzina and Zougari, 2008; Alhammadi and Edward 2009; Ezz et al., 2010; Marzouk 2011). This confirmed our first hypothesis also for other elements. Environmental factors, such as the availability of nutrients in the soil, time in the growth season, position of the leaf sampled on the tree, part of the leaf sampled, and age of the plants affect element composition of any plant. Therefore, it is not surprising that the concentrations of essential elements measured in Experiment 1 were often different than those determined by the Critical Value Approach, which describes the optimal elemental concentration (Labaied et al., 2020). The average translocation factors (across the varieties) suggest that Mg, S, Fe and Cu were retained in roots, whilst P, Ca, Mn and Zn, were accumulated in leaves (**Table S2**). There are few studies where both roots and leaves were analysed for their micronutrient composition. At least for Ca, observations from the present study are consistent with previous data (Alhammadi and Edward, 2009). The general observations that concentrations of Ca and Mn in leaves are greater than in roots, and that Fe and Cu are predominantly partitioned into roots, are consistent with angiosperm species from different orders, whereas the partitioning of other elements including Mg cannot generally be observed in angiosperms from other clades (Neugebauer et al., 2020).

Due to obvious differences in the ability to restrict Na accumulation in leaves between Sultana and other date palm varieties, we continued detailed analysis on this variety. For comparison we selected the variety Khalas. It has been the subject of several previous studies, therefore allowing for data comparison (Radwan et al., 2015; Al-Qurainy et al., 2017; Yaish et al., 2017; Mueller et al., 2023).

### Differences in Na Translocation Between Khalas and Sultana can be Explained by Different Na Allocation Mechanism

Varieties Khalas and Sultana were grown in parallel in Experiment 2. At harvest, root tips and the root bases were excised from roots and shoot bases, leaf bases and leaf tips were excised from shoots. Cross-sections were prepared by cryo-sectioning and freeze-drying (**Figure S7**). The remaining material was used to determine bulk element concentrations. Tissue concentrations of Na and Cl were larger in Experiment 2 (**Figure S8A, B**) compared to Experiment 1, which is a consequence of somewhat larger NaCl supply. However, the K concentrations were comparable with data from Experiment 1 (**Figure S8C**). Translocation factor for Na in Sultana was smaller than in Experiment 1 but was still larger at NaCl treatment than in Khalas (**Table S3**), which indicates stability in the differences in Na translocation factors under NaCl in these two varieties.

The spatial distribution of elements was determined in cross-sections of the sampled five plant parts, by multielement and quantitative micro-PIXE with 4-5 µm lateral resolution. There were no obvious differences in the location of Na, Cl and K between the two varieties (**Figure 4 and Figure S9**) and no indication that Na replaced K as an osmoticum. In plants that had not been exposed to NaCl tissue Na concentrations were below the detection limit of the micro-PIXE technique. The ability to detect Na distributions declined in the direction of the transpiration stream (**Figure 4 and Figure S9**) in agreement with the decrease in Na concentrations from root tip to shoot tip (**Figure S8**). The largest Na concentrations in root tips for Khalas variety were previously reported by Mueller et al., (2023). In root tips, most Na was present in the cortex cells (**Figure 4 and Figure S9**). At the base of the roots, the rhizodermis contained Na-rich areas which were only observed in the cross-section of Khalas, but not in Sultana.

**Figure 4.**
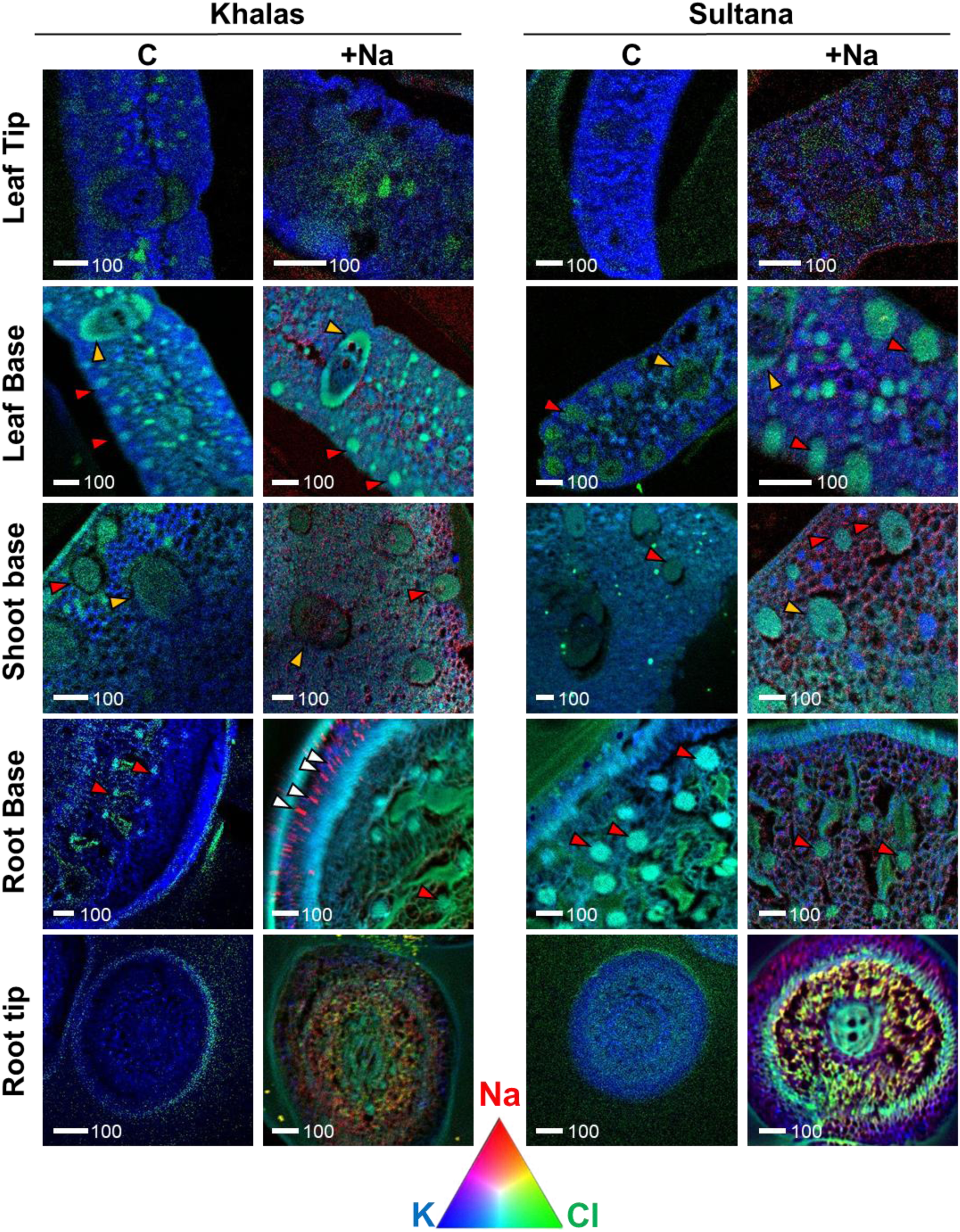
Overlay of representative tissue-specific distribution maps of sodium (Na) in red, chlorine (Cl) in green and potassium (K) in blue, in different parts of two date palm (*Phoenix dactylifera*) varieties. Plants were irrigated with double-distilled water (Control, C) or with 150 mM NaCl for two weeks and 300 mM NaCl for six weeks (+Na); Experiment 2. Scale bars represent 100 µm.

The ability of Khalas to form these Na hotspots, presumably immobilising Na, may explain poorer mobility of Na in Khalas compared to Sultana (**Figure 2**), which confirms our second hypothesis that differences in Na translocation can be explained by different Na allocation mechanism(s), which was true for the root Na allocation. Large Na accumulation in epidermis was observed in moderately salt tolerant barley (*Hordeum vulgare* L.), which had large Na concentration in the stellar tissues too (Munns and Tester, 2008). Similarly, in roots of halophyte *Aster tripolium* L., a decrease in Na concentration from rhizodermis to the stele was reported while Cl was distributed evenly (Scheloske et al., 2004). Similar Na distribution in roots was observed in a desert halophyte *Bassia indica* (Wight) A. J. Scott (Shelef et al., 2016). Thus, date palms seem to restrict Na transport to the vasculature, in an effort of retaining it in roots, as a general Na tolerance mechanism. There was no apparent co-localisation of K or Cl with Na in roots (**Figure 4**), but there was a marked correlation between K and Cl with Na in distribution images of root tips and root bases of Sultana, which was lacking in Khalas (**Figure S10**).

Leaves are believed to be even more sensitive to Na toxicity, therefore, Na exclusion systems (e.g. salt glands) can be useful to decrease Na concentration in leaves in some plant species (Dassanayake and Larkin 2017). Salt glands have not been reported for date palms. Therefore, dealing with NaCl/osmotic stress in leaves is the only option and it is most often accomplished by compartmentation at tissue and cell levels. For example, allocation of Na away from photosynthetic tissues has been demonstrated in leaves of *B. indica*, which have large water storage tissues where Na was accumulated and presumably enabled large K/Na quotient in mesophyll compared to other leaf tissues (Pongrac et al., 2013). In stems and leaves of date palms Na was evenly distributed, thus tissue specific Na compartmentalisation does not appear to occur. Although larger K concentrations in distribution maps of the leaf base and in leaf tips were observed in Sultana than in Khalas indicating a more favourable K/Na quotient for these regions (**Figure S10**).

In roots, in shoot stem, and at the base of the leaf, Cl was allocated to fibre bands peripherally positioned in the leaf and large sclerenchyma sheaths of vascular bundles (**Figure 4 and Figure S9**). This was a peculiar observation as Cl acts as a major osmotically active solute in the vacuole and is involved in both turgor- and osmoregulation (White and Broadley 2001), therefore vacuolar compartmentalisation was expected. The fibre bands and the sclerenchyma sheaths of vascular bundles were accompanied by stegmata, Si accumulating cells (**Figure S11**) as reported previously (Bokor et al., 2019; Midani et al., 2020). Sulphur was found predominantly in epidermal and sub-epidermal tissues, P in the vasculature, and Ca in mesophyll cells in leaves and in hotspots in roots (**Figure S11**). There is no clear explanation for the Cl-rich fibre bands in date palms which in planta serve to mechanically strengthen the leaves and have important technological value (Midani et al., 2020).

### Differences in Na Translocation in Khalas and Sultana are Due to Different Expression Pattern of NaCl-Responsive Genes

When comparing gene expression of the two varieties, with contrasting abilities to restrict Na accumulation in leaves, many gene expression differences were found already under control conditions (**Figure 5**).

**Figure 5.**
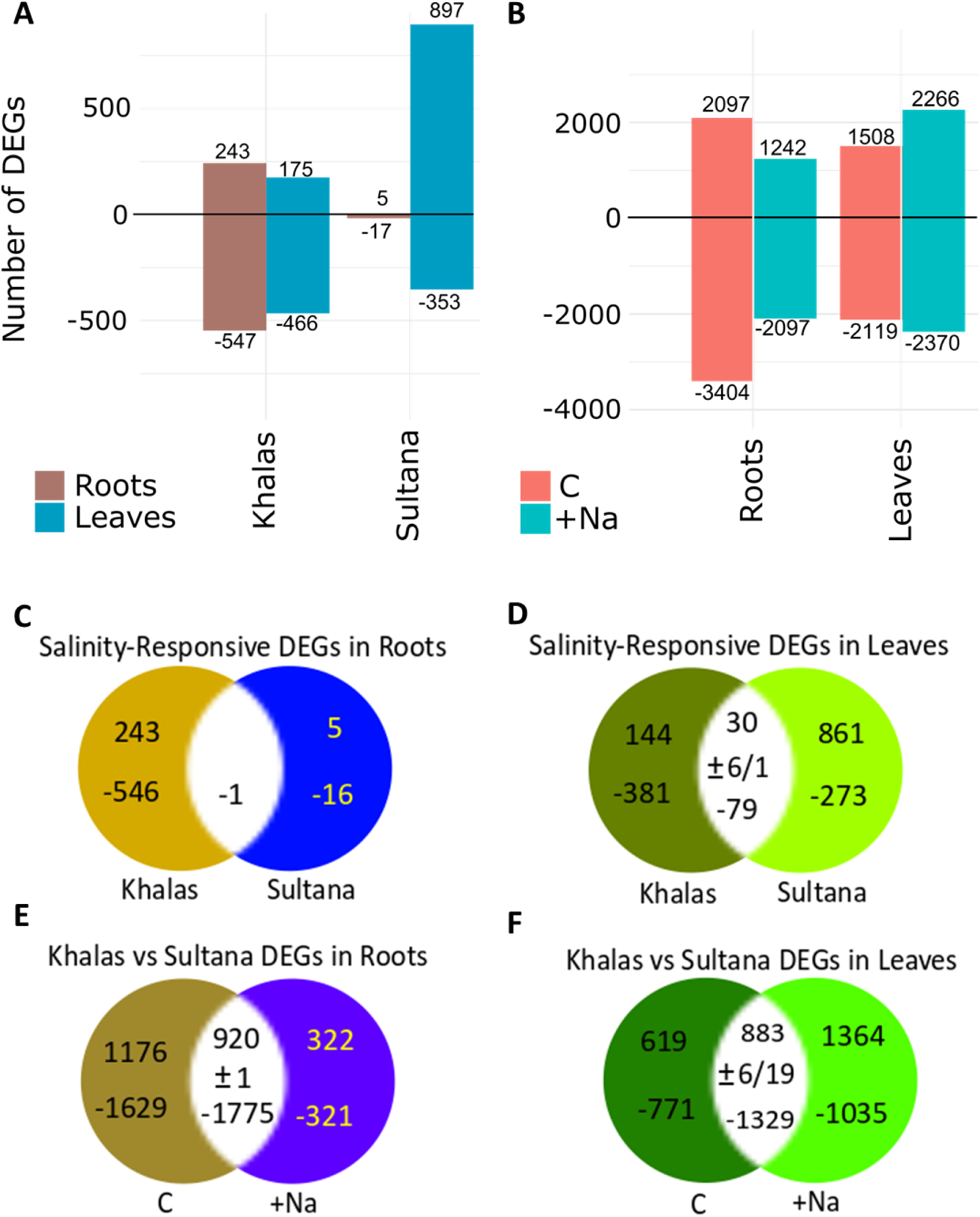
Number of differentially expressed genes (DEGs; A, B) and Venn analysis of DEGs (C-F) in different tissues of two date palm (*Phoenix dactylifera*) varieties. Differentially expressed genes were either induced (+) or repressed (-) in saline conditions (+Na; watered with NaCl for two weeks with 150 mM and for six weeks with 300 mM) compared to non-saline conditions (C; watered with double-distilled water only) in roots (C) or in leaves (D) or in the two varieites in C or +Na treatments in roots (E) or leaves (F); Experiment 3.

This extraordinary difference in gene expression under control conditions indicates that they are very different genetically, highlighting their potential for different performance under NaCl stress. This extensive genetic difference, which is not expressed at phenotype level or does not affect the fitness of a plant, may be a result of neutral molecular variation (Chung et al., 2023). Most NaCl-responsive genes were unique to either variety and hardly any common gene ontology (GO) enrichments were found between Khalas and Sultana (**Figure 6**). This indicates a substantially different gene regulation in general followed by a drastically different response to NaCl. It highlights several genes which could be responsible for increased NaCl tolerance in Khalas and helps to understand the variation in NaCl detoxification strategies in date palms.

**Figure 6.**
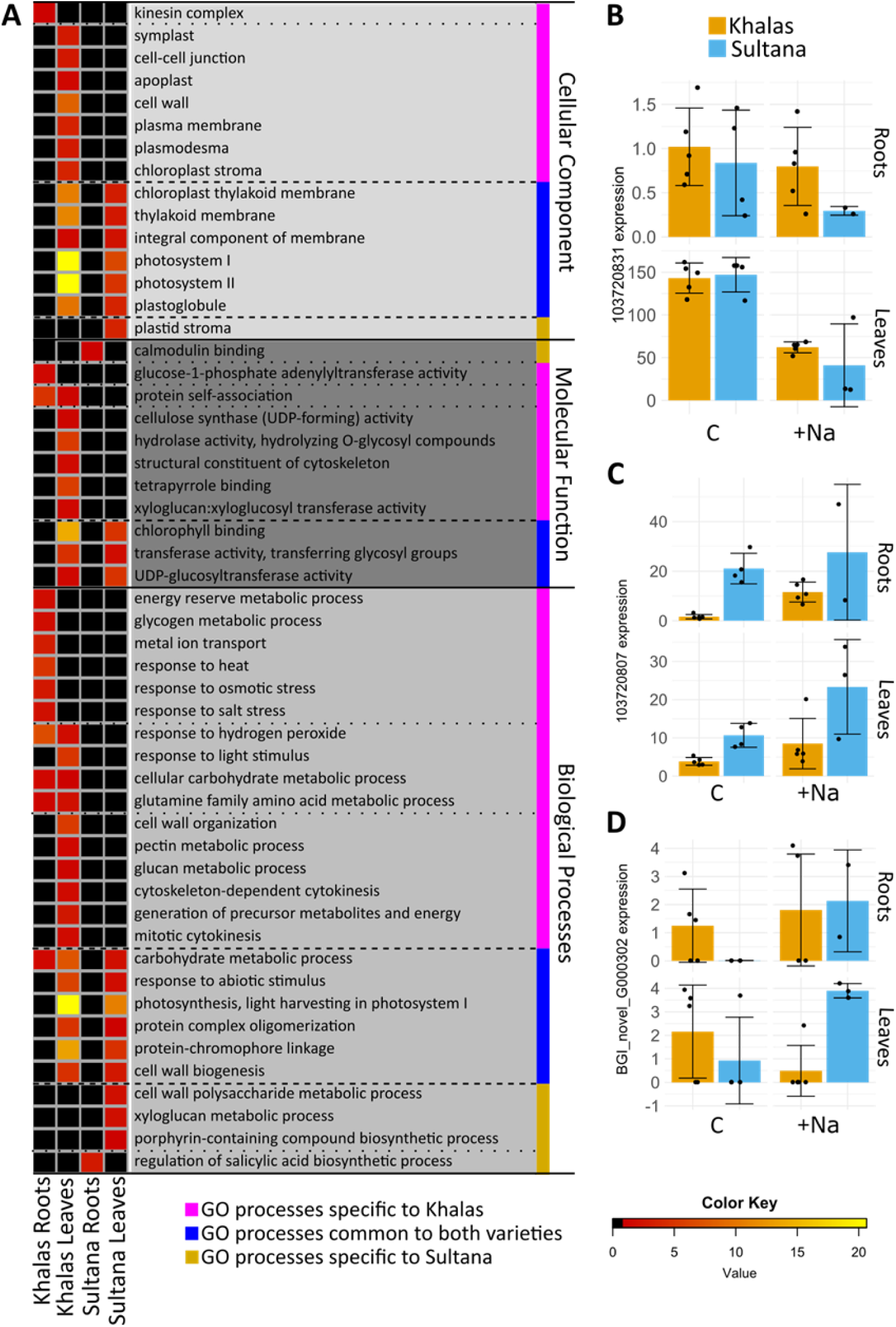
Gene ontology (GO) enrichment heatmap with p values for GO groups enriched among differentially expressed genes (DEGs) in two date palm (*Phoenix dactylifera*) varieties. Plants were exposed to salinity (+Na; watered with NaCl for two weeks with 150 mM and for six weeks with 300 mM) and to non-saline conditions (C; watered with double-distilled water only). GO were grouped into three categories (shaded in different greys) and within each further sorted according to their occurrence in Khalas (purple), in both varieties (blue) or in Sultana (gold) (A). Examples of gene expressions of genes belonging to GO groups: from the GO „Photosysthem I“, a GO group overrepresented among DEGs in both Khalas and Sultana leaves (B), from the GO „Response to salt stress“, a GO group overrepresented amond DEGs only in roots of Khalas (C), and the GO group „Regulation of SA biosynthetic process“ overrepresented among DEGs in roots of Sultana (D).

Differentially expressed genes (DEGs) were defined between treatments (NaCl treatment to Control) and between varieties for each organ (roots and leaves). All DEGs can be found in **Tables S4-11** available at https://doi.org/10.5281/zenodo.14283064. The expression of 790 genes was altered by NaCl in roots of Khalas, but only of 22 genes in roots of Sultana (**Figure 5**). In leaves the expression of 641 genes of Khalas and 1250 genes of Sultana was altered by NaCl (**Figure 5A**). The DEGs between varieties show differences in gene expression due to the genetic background (**Figure 5B**). This reveals larger number of DEGs between varieties than due to NaCl. Finally, by comparing NaCl-responsive genes (**Figure 5A**) between cultivars (**Figure 5C and D**) we show commonalities and differences in their response to NaCl. In roots fewer common DEGs were seen compared to leaves. These observations allow us to conclude that these two varieties respond drastically differently to NaCl on a gene expression level and therefore supports our third hypothesis that Na translocation in Khalas and Sultana is due to different expression pattern of NaCl-responsive genes. Below we survey the DEGs and the corresposnding GO identified and relate them to salinity, when possible, by comparing them with current understanding of reponse to NaCl in *Arabidopsis thaliana* (annotation of which was used to derive gene functions).

### NaCl-Responsive Genes in Roots of Khalas, but not of Sultana, were Linked to Growth and Heat and Salt Stress

In roots, only one gene was NaCl-responsive in both varieties, while there were 116 such DEGs in leaves (**Figure 5C, D**). Among these up and down regulation was conserved, except for seven genes in leaves which were upregulated in one variety but downregulated in the other (**Figure 5D**). The lack of a transcriptomic response to NaCl in roots of Sultana, as compared to Khalas, may reflect its apparent inability to limit Na mobility (**Figure 2**). Contrastingly, the large number of genes responding to NaCl in shoots of Sultana (**Figure 5A**) may represent the greater metabolic adaptation due to relatively large Na concentration in leaves (**Figure 1A**). This observation is also supported in a Principal Component Analysis (**Figure S12)**. A comparison of DEG and pathways of differentially expressed genes between Sultana and Khalas may identify candidate genes responsible for the differences in Na translocation factors and NaCl tolerance observed.

Gene ontology enrichments were studied among DEGs (**Figure 6**). This analysis detects pathways which occur less or more often than would be expected by chance. If a pathway is relevant to an environmental stimulus or to a genotype we expect several of its components to be changed. In a summary of the GO enrichment analysis, groups of processes which were changed in either one or both varieties were displayed (**Figure 6A**). In Khalas roots for example, expression of genes related to kinesin complex was significantly changed (**Figure 6A**). Eight genes with this GO annotation were down regulated (**Table S4-7**). Kinesin motor proteins transport cargo along microtubules and are involved in various cellular processes such as organ organisation or vesicle transport and ultimately cell growth. Thus, their change may be a response to salinity by changes in growth to avoid further Na exposure (Pierik and Testerink 2014). However, root biomass of Khalas in Experiment 1 was not affected by salinity (**Figure S1**) and in Experiment 3 biomass was not determined due to minimal plant handling required before plant processing. Further metabolic changes observed in Khalas roots were connected to energy metabolism and to 16 known stress response genes (**Figure 6A**) all except one of which were induced under NaCl treatment. Of these 16, 11 were heat shock proteins and two are heat shock transcription factors. There are many examples which show a cross- talk between heat and salinity stress and various authors reported an increased expression of heat shock proteins to lead to salinity tolerance (Song and Ahn 2011; Zang et al., 2019; Al Khateeb et al., 2020). Additionally, previous RNAseq studies on date palm have identified induction of heat shock proteins after salinity treatments (Yaish et al., 2017; Mueller et al., 2023). Also, among the changes specific to Khalas roots is a response to salt stress. The expression of one gene belonging to this group shows the induction in Khalas roots (**Figure 6C**). This GO group is directly linked to known NaCl tolerance mechanisms. Furthermore, changes in the expression of metal transporters seen in Khalas roots (**Figure 6A**) may have limited substrate specificity or diverging substrate specificity and as such be involved in restriction of Na mobility within this variety. Five DEGs were metal ion transmembrane transporters and could be directly involved in maintaining a sufficient K/Na quotient (LOC103713167, probable potassium transporter 11 was induced by salinity; LOC120109241, two-pore potassium channel 5-like was repressed; LOC103712998, metal tolerance protein 4- like was repressed; LOC103722218, probable magnesium transporter NIPA4 was repressed; LOC103720046 - two-pore potassium channel 5-like was repressed; **Table S4**).

The molecular function “glucose-1-phosphate adenyltransferase activity” were also differentially regulated only in roots of Khalas (**Figure 6A**). One of the three DEGs were repressed while two were induced (**Table S4-7**). This GO category is important in transient and storage starch synthesis (Figueroa et al., 2022).

### NaCl-Responsive Genes in Shoots Both Varieties were Linked to Photosynthesis and Cell Wall Growth Indicating Stress Response

Several GO groups were enriched among DEGs of both Khalas and Sultana leaves, but none were common in roots (**Figure 6A)**. The locations mostly targeted for transcriptomic changes in both varieties were related to photosynthesis: thylakoids and photosystems. They were repressed in leaves due to NaCl in both varieties (**Figure 6B**). This observation is in line with previous evidence showing a decreased photosynthetic activity in salt-sressed date palms (Ramoliya and Pandey 2003; Sperling et al., 2014; Yaish and Kumar, 2015; Al Kharusi et al., 2017; Du et al., 2021). Additionally, cell wall biogenesis genes were changed by salinity. Interestingly, only three cell wall genes were depressed in both varieties (**Table S4-7**). Additionally, in Sultana 17 distinct cell wall biogenesis genes and Khalas 14 other genes were differently expressed (**Table S12**). While Khalas repressed 12 out of 17 genes, Sultana induced 13 and repressed only 7 indicating different strategies to ameliorate NaCl stress through the cell wall processes. Among the strongly induced genes was LOC103702579: cellulose synthase-like protein E6 (**Table S7**). Cellulose synthase is required for cell wall production and therefore cellular growth processes (Festucci-Buselli et al., 2007).

Certain GO groups were enriched specifically in Sultana leaves, namely “plastid stroma” or “cell wall polysaccharide metabolic process” (**Figure 6A**). In Sultana roots, two genes of the GO group “regulation of salicylic acid biosynthetic process” were up (**Figure 6D**) and down regulated, respectively. Their function is unknown and thus it is unclear if they enhance salicylic acid (SA) synthesis or repress it. Salicylic acid is involved in a number of processes within plants and has been shown to regulate a plants response to salinity (Hassoon and Abdulsattar Abduljabbar, 2020). In date palms SA treatment was shown to improve growth, chlorophyll content, free radical damage, water content and electrolyte leakage (Shareef et al., 2022). We could thus speculate that Sultana, unlike Khalas, utilises SA-based response to ameliorate effects of NaCl.

Few commonalities with GO enrichments detected in our study were found from a recent study in Khalas by Mueller et al., (2023). One possible reason is that authors combined both gene expression and protein studies in their network analysis, while only transcriptomics were provided in our study. One identifiable exception was a GO group “active ion transmembrane transport” and “metal ion transport”, which were overrepresented among the DEGs in both studies. Mueller et al., (2023) observed a correlation with the gradient in Na concentration from the tip to the base of the root and increased expression of one of three *SOS1* genes. The *SOS1* (Salt Overly Sensitive1) genes code for plasma membrane Na/H antiporters and have been shown to protect *A. thaliana* from salinity stress (Xie et al., 2022). A *sos1* mutant shows sensitivity to external Na and increased shoot Na concentrations (Shi et al., 2002). It has been shown that SOS1 function depends on its tissue-specific location: SOS1 located in the outer root tissues exports Na from root tissues while SOS1 located at the stele is relevant for shoot directed Na transport (Munns and Tester, 2008). It was recently proposed, that SOS1 activity at the stele leads to enhanced water transport, using Na as an osmoticum (Foster and Miklavcic 2019). Thus, high expression of SOS1 in date palm roots could indicate an exclusion strategy as well as an osmotic adjustment strategy depending on the location of the expression. Increased *SOS1* expression may also be a result of increased Na concentrations as the gene is responsive to NaCl treatment (Shi et al., 2002). In Khalas shoots two potential SOS1 orthologues have been reported (Mueller et al., 2023), but none were found to be differentially expressed in Khalas in our study. By contrast, an up regulation of LOC103713094, sodium/hydrogen exchanger 8 (**Table S7**), the closest orthologue of AtSOS1 according to our BLAST analysis was found in Sultana shoots. Little is known about the function of SOS1 in shoots as most studies in salinity tolerance have focused on roots. Nevertheless, few studies have shown SOS1 activity to be regulated in liaison with different genes in shoots and roots (Yin et al., 2020) but there is little evidence to show how the gene functions to alleviate salinity stress. As an exporter, it may function to unload Na from the xylem or to move excess Na into the apoplast. Induction in Sultana could indicate the latter, as a compartmentalisation strategy. In *A. thaliana* the SOS pathway is well described and important for salinity tolerance (Quintero et al., 2002). SOS2, a protein kinase, and SOS3, a Ca^2+^ sensor, are as important as their target SOS1 for Na tolerance. Neither variety exhibited a change in expression of LOC103708740, CBL-interacting protein kinase 24-like, orthologue of SOS2 or LOC 103711181, calcineurin B-like protein 7, orthologue of SOS3.

### Involvement of Transporters in NaCl-Response

Another mechanism for NaCl tolerance is its sequestration in vacuoles. In *A. thaliana* this is performed via NHX1, a vacuolar Na uptake transporter (Gaxiola et al., 1999). In roots of Khalas the expression of PdNHX1 was not affected by salt stress analysis while in leaves the gene was down regulated (Mueller et al., 2023). A LOC103723229, sodium/hydrogen exchanger 2-like isoform X1, a close BLAST match for NHX1 was found to be induced in shoots of Sultana in our study, suggesting preferentially storage of Na in leaf vacuoles as a detoxification strategy. This observation is in line with previous studies of the transporter in *A. thaliana* which showed induction of the gene after salinity treatment (Yokoi et al., 2002).

A further transporter believed to be related to salinity tolerance is HKT1 (Rus et al., 2001, 2004). It is part of the high-affinity K^+^ transporter (HKT) transporter/channel family (Horie et al., 2009). The protein is preferentially transporting Na and responsible for Na distribution within plants (Mäser et al., 2002). In *A. thaliana* the transporter has very specific functions, which depend on the localisation (An et al., 2017), sequence (Baxter et al., 2010) and expression strength (Rus et al., 2006). A hypofunctional allele of HKT1 in accessions located close to the coast like Ts-1 accumulate large amounts of Na in their leaves (Rus et al., 2006) but is hyper-functional in flower and reduces Na concentrations locally, thus maintaining seed production (An et al., 2017). Three *HKT1* orthologues were repressed by salinity in roots and one in shoots of Khalas (Mueller et al., 2023) and down regulation of HKT1 after a short burst of Na-treatment in Khalas roots has been observed (Yaish et al., 2017). It is possible that in this variety HKT1 function is linked to Na uptake from the soil and the gene therefore down regulated at high external Na to restrict uptake. Or, similarly to *A. thaliana* accessions growing close to coastal regions, the gene may have different expression patterns in other organs, effectively enabling it to restrict Na to less sensitive tissues (An et al., 2017). Down regulation of HKT1 orthologue LOC103701574, cation transporter HKT8-like and LOC103701576, probable cation transporter HKT6, most similar to AtHKT1 in a BLAST search, was observed in shoots of Khalas (**Table S5**). By contrast, in Sultana shoots an induction of LOC103715558, cation transporter HKT1-like, an *AtHKT1* orthologue according to BLAST was found (**Table S7**). We conclude that Khalas and Sultana use HKT1 very differently. It could for example, be used to export Na from the root if located in the epidermis. Further insight into the special resolution of these transporters, and importantly their variants, could provide insight into Na detoxification mechanisms.

Vacuolar compartmentalisation through NHX2 has been reported as a Na tolerance strategy. Two orthologues of the vacuolar NHX2 transporter were induced in roots of Khalas (Mueller et al., 2023). By contrast, no NHX-like genes were detected among the DEG after salinity treatment in Khalas and Sultana, but a number of ATPases, many plasma membrane types among them, with the DEGs in shoots in Sultana and Khalas and roots of Khalas (**Table S7, S5 and S4,** respectively) were found. These transporters are important to establishing electrochemical gradients, also between vacuole and cytosol, and thereby contribute to ion transport and compartmentalisation. Such induction of vacuolar ATPases have been observed in Khalas before (Yaish et al., 2017).

An important group of metal transporters for salinity tolerance are the KUP/HAK/KT family (Mäser et al., 2001). Maintaining K homeostasis is crucial under salinity stress and many members of this family are, despite being primarily K transporters, accessible to Na. In roots of Khalas, two DEGs of the KUP/HAK/KT family were (**Table S4**, one was induced while the other was repressed by NaCl) while none were regulated by NaCl in Sultana (**Table S6**). In shoots, several genes in both varieties showed differential expression. A highly expressed gene LOC103713167, KUP system K uptake protein, was induced by NaCl in roots of Khalas (**Table S4**). Potassium concentrations in leaves were not reduced by NaCl treatment (**Figure 3**). Khalas may induce the K transporter in response to NaCl stress to maintain K uptake. The most relevant K uptake transporters in *A. thaliana* HAK5 (LOC103712364) and AKT1 (LOC 103721181) were not differentially expressed and were induced in shoots of Sultana only. In roots on the other hand a LOC103713167, probable K transporter 11 was induced and LOC103706783, KUP system K uptake protein and LOC103702633, K channel AKT2 were down regulated. Consistent with previous reports, this could indicate an intensified K uptake or a mechanism to avoid Na uptake through unspecific action of K transporters (Yaish et al., 2017; Mueller et al., 2023). Many K transporters were differentially expressed between roots of Khalas and Sultana roots (**Table S4 and 6**). 25 were repressed and 13 up regulated in Sultana. In shoots likewise there were many differences, 14 genes were repressed and 10 induced.

### NaCl-Responsive Building of the Translocation Barrier in Roots

The Casparian strip is an endodermal structure with important role in preventing the non-selective apoplastic bypass of salts into the stele along the apoplast (Chen et al., 2011). We noted that LOC103720750, Casparian strip membrane protein 1-like, an orthologue of *AtCASP4*, was repressed in roots of Khalas but not of Sultana (**Table S4 and S6, respectively**). This tallies with Khalas’s abilities in modulating responses to NaCl in roots and retaining a small Na translocation factor. Modulations in the Casparian strip in response to NaCl stress have been observed in *A. thaliana* (Wei et al., 2021). The formation of the apoplastic barrier strongly correlates with the shoot ion accumulation of K (positively) and Na (negatively) (Reyt et al., 2020). Previous studies have shown upregulation of *PdCASP5* (Yaish et al., 2017) and a positive impact of *SbCASP4* on salt tolerance (Reyt et al., 2020). Thus, the role of individual CASP proteins may be different in date palms and will require further studies.

### Differentially Expressed Genes Related to Compatible Solutes and Antioxidants

One acclimatory mechanism enabling plants to cope with the osmotic stresses associated with salinity is the accumulation of compatible solutes, such as sugars, sugar alcohols, quaternary ammonium compounds and amino acids (Slama et al., 2015). Previous studies characterising the metabolic responses of date palm suggest that the concentrations of sugars and sugar alcohols in roots and leaves often remain unchanged or decrease when plants are exposed to NaCl (Mueller et al., 2023), although there are contrary reports (Al Kharusi et al., 2019b). We saw an overrepresentation of xyloglucan metabolic processes in shoots of Sultana, hinting that sugar compounds may be altered in response to NaCl. Several sugar transporters were also down regulated in both Khalas and Sultana as was seen before for Khalas (Mueller et al., 2023).

Nitrogen compounds (amino acids proline, phenylalanine, ethanolamine leucine, isoleucine, γ- aminobutyric acid (GABA) and butyro-1,4-lactam) have been shown to accumulate in roots and leaves of date palm exposed to salinity (Yaish 2015; Al-Qurainy et al., 2017, 2020; Jana et al., 2019; Al Kharusi et al., 2019b; Du et al., 2021; Mueller et al., 2023). Likewise, overrepresentation of glutamine family amino acid metabolic processes in roots and shoots of Khalas was observed (**Figure 6A**). This variety probably uses glutamine to ameliorate toxic effects of NaCl. Furthermore, amino acid transporter was also among the DEGs in Khalas. Notably the only commonly downregulated gene in roots of Khalas and Sultana and in a previous study of Khalas was LOC103718051, a transmembrane transporter and orthologue to *A. thaliana UMAMIT33* (*Usually Multiple Acids Move In and out Transporter 33*). It could be involved in amino acid exporter although the gene itself has not yet been characterised (Besnard et al., 2018).

Antioxidants are important in countering oxidative stress, which increases in response to NaCl stress (Al Kharusi et al., 2019a). Correspondingly, a response to hydrogen peroxide processes was overrepresented in Khalas but not Sultana (**Figure 6A**). Salinity stress generally increases the concentrations of superoxide dismutase (SOD), catalase (CAT) and ascorbate peroxidase (APX) in roots and leaves of date palm varieties that are relatively tolerant to soil salinity (Abdulwahid, 2012; Darwesh, 2013; Al-Qurainy et al., 2017, 2020; Al Kharusi et al., 2019a). Several catalase and peroxidase genes were regulated in roots and shoots of Khalas and shoots of Sultana as seen before (Yaish et al., 2017; Mueller et al., 2023). Consequently, antioxidants often accumulate in roots and leaves of NaCl-stressed date palms (Al Kharusi et al., 2019b, 2021).

## Conclusions

Sodium and Cl accumulation across six date palm varieties was evaluated. No growth penalties were observed due to NaCl treatment, but a significant difference in element partitioning between varieties and particularly Na translocation behaviour for two varieties: variety Sultana had the largest Na translocation factor, while the variety Khalas exhibited better restriction of Na mobility. These two varieties were studied in greater detail. There were no apparent differences in element distribution across tissues between the two varieties, except at the root base, where Na hotspots were observed in Khalas but not in Sultana indicating that Khalas restricts Na translocation to shoot by immobilising Na at the root base. Sodium accumulated in cortex in root tips and was evenly distributed in leaves, while Cl accumulated in fibre bands and sclerenchyma sheaths of vascular bundles. There were dramatic differences in gene expression between varieties and between organs in both control and NaCl treatments. In general, Khalas responded mainly in roots and Sultana in leaves, which supports our hypothesis that there is sufficient differences in the expression pattern of NaCl-responsive genes to allow for the phenotypic differences observed. Differentially expressed genes in roots of Khalas were linked with root growth and modulation of the Casparian strip, stress responses and metal transporters, which presumably limit Na movement to the leaves. Sultana on the other hand modulated growth and cell wall processes in the shoot while using silicic acid to signal NaCl stress in roots.

## Author Contribution

PJW conceived and managed the project, designed the experiment and wrote the first draft. GW and JAT performed and analysed all plant material from Experiment 1. PP performed Experiments 2 and 3, prepared samples and performed localisation analyses from Experiment 2 and wrote the manuscript. SF prepared samples and managed the RNAseq for Experiment 3 and wrote the manuscript. MK technically supported element localisation analysis at the nuclear microprobe. KN and LKB performed element analyses for Experiment 2. KVM financially contributed to supporting Experiments 2 and 3 and RNAseq, analysed root sections to confirm initial observation and contributed to the final version of the manuscript. MŠ analysed root sections to confirm initial observation and edited the final version of the manuscript. HAE financially contributed to supporting Experiment 1. PP and SF contributed equally.

## Acknowledgements

We thank Martin R. Broadley for comments on the manuscript. This paper is dedicated to the dear memory of Philip J. White, who sadly passed away during the final stages of drafting this manuscript.

## Data Availability Statement

Majority of the data that support the findings of this study are available in the Supplementary file and RNAseg dataset is available at https://doi.org/10.5281/zenodo.14283064. Everything else will be available from the corresponding author upon reasonable request.

