## Supplementary files for "Element partitioning, element localisation and transcriptome responses of date palms exposed to NaCl"

**Running title:** Contrasting root Na location and transcriptome responses of two date palm varieties

***Corresponding author:**

Paula Pongrac, University of Ljubljana, Biotechnical Faculty, Jamnikarjeva 101, Ljubljana, Slovenia

[¥]These authors contributed equally

[‡] Current position: Albrecht Daniel Thaer-Institute of Agricultural and Horticultural Sciences, Faculty of Life Sciences, Berlin Humboldt University, Invalidenstraße 42, 10115 Berlin, Germany

Number of Supplementary Figures: **12**

Number of Supplementary Tables: **3** (Tables S1-S3)

Number of Supplementary Tables available at https://doi.org/10.5281/zenodo.14283064: **9** (Tables S4-S12)

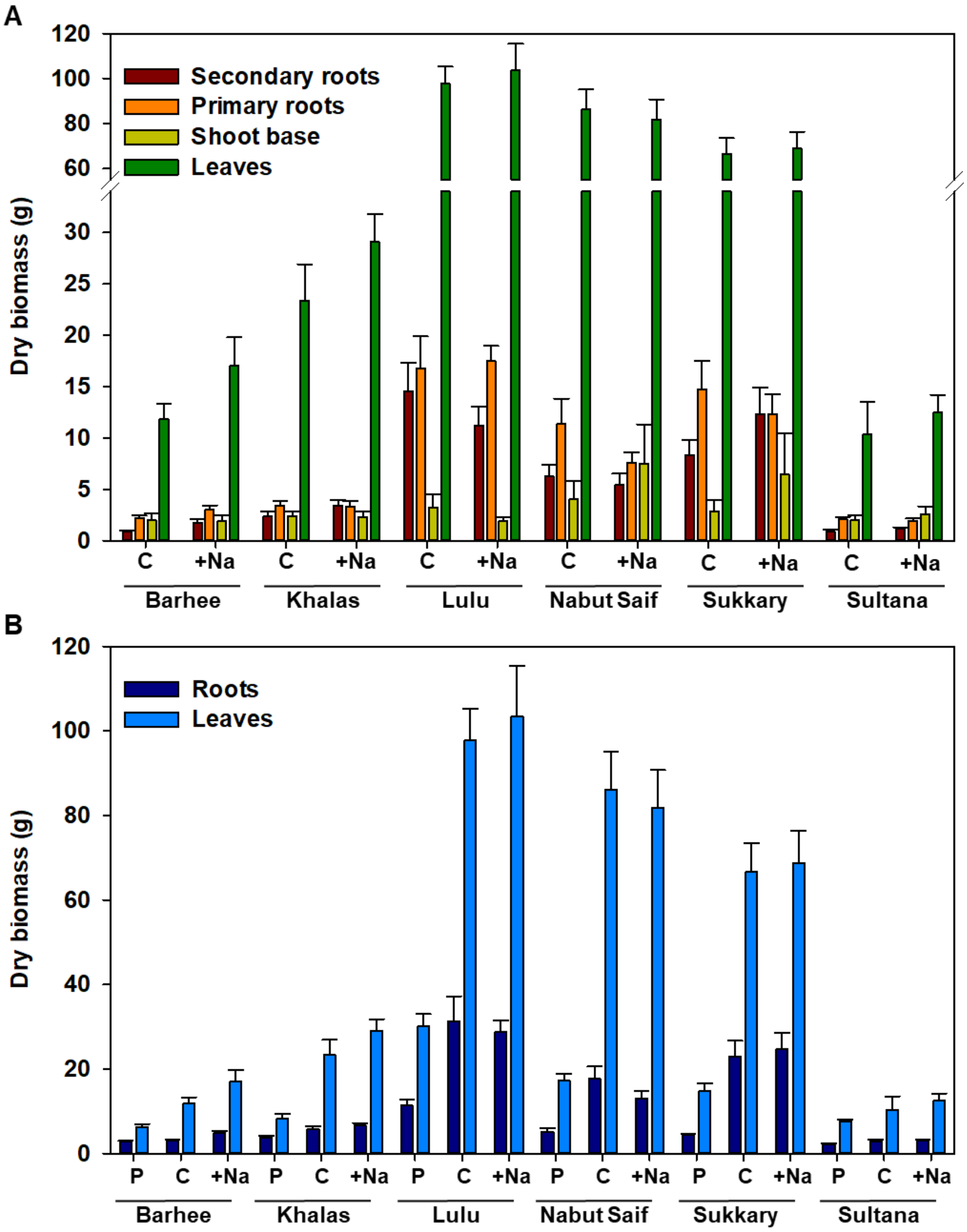


**Figure S1** Dry biomass of different parts (A) and of roots (secondary plus primary roots) and leaves (B) of six date palm (*Phoenix dactylifera*) varieties. Plants were analysed immediately after purchasing (pre-experiment plants; P), irrigated with double-distilled water (Control, C) or once with 400 mL of 150 mM NaCl at the beginning of the experiment, then after two, four and six weeks with 400 mL of 300 mM NaCl (+Na); Experiment 1. Shown are averages ± standard errors (n=6). DW, dry weight.

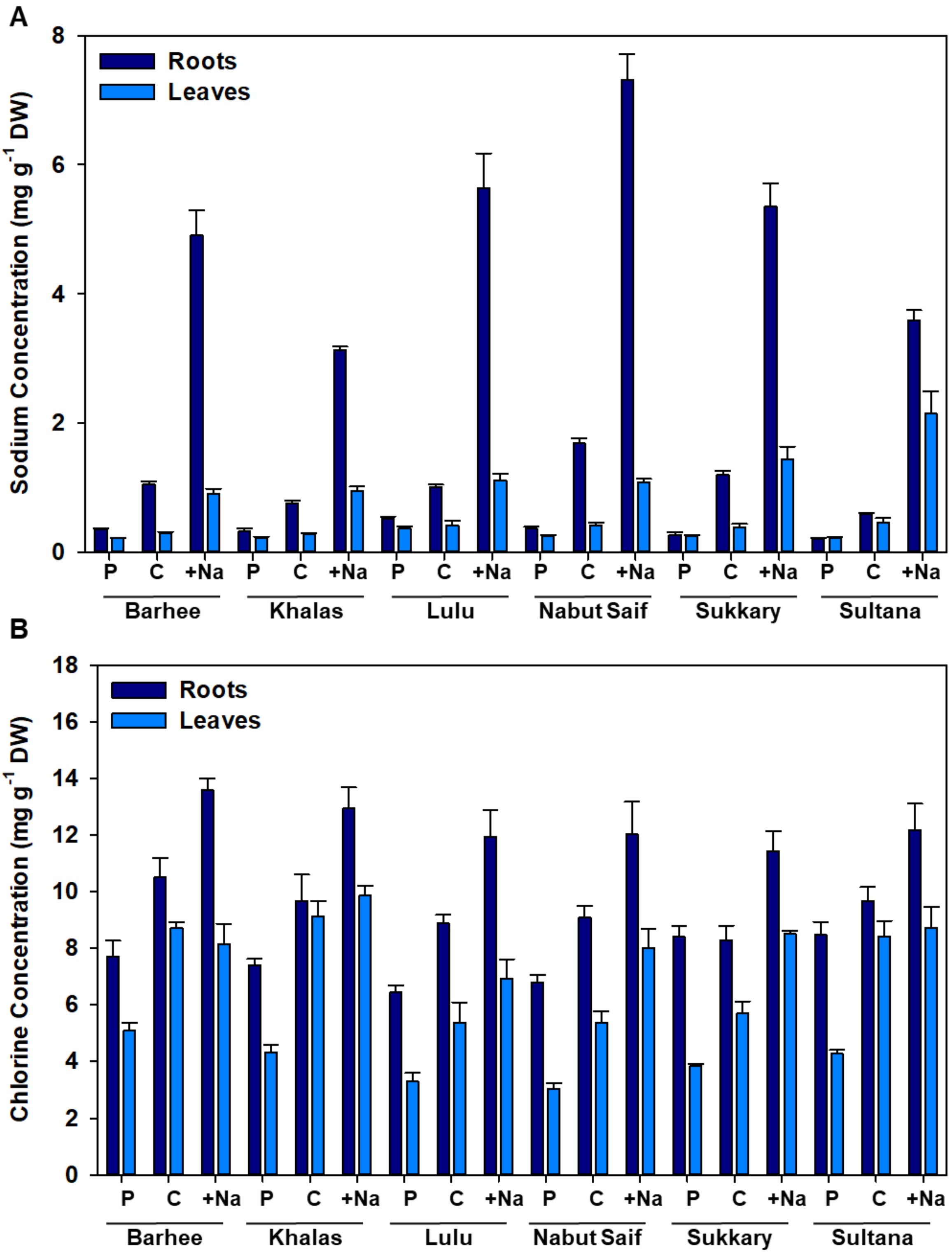


**Figure S2** Sodium (A) and chlorine (B) concentration in roots and leaves of six date palm (*Phoenix dactylifera*) varieties. Plants were analysed immediately after purchasing (pre-experiment plants; P), irrigated with double-distilled water (Control, C) or once with 400 mL of 150 mM NaCl at the beginning of the experiment, then after two, four and six weeks with 400 mL of 300 mM NaCl (+Na); Experiment 1. Shown are averages ± standard errors (n=6). DW, dry weight.

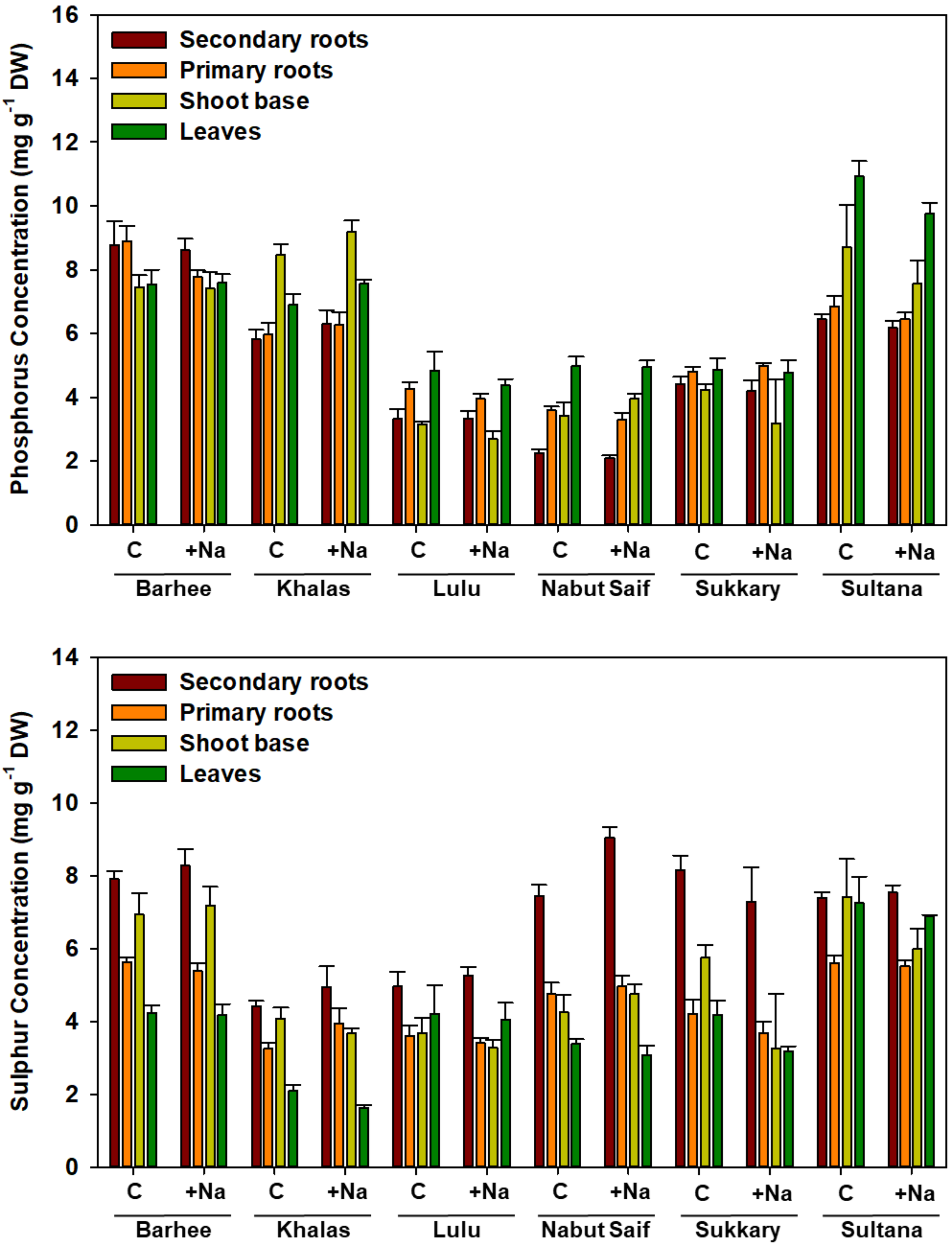


**Figure S3** Phosphorus and sulphur concentration in different parts of six date palm (*Phoenix dactylifera*) varieties. Plants were irrigated with double-distilled water (Control, C) or once with 400 mL of 150 mM NaCl at the beginning of the experiment, then after two, four and six weeks with 400 mL of 300 mM NaCl (+Na +); Experiment 1. Shown are averages ± standard errors (n=2-6). DW, dry weight.

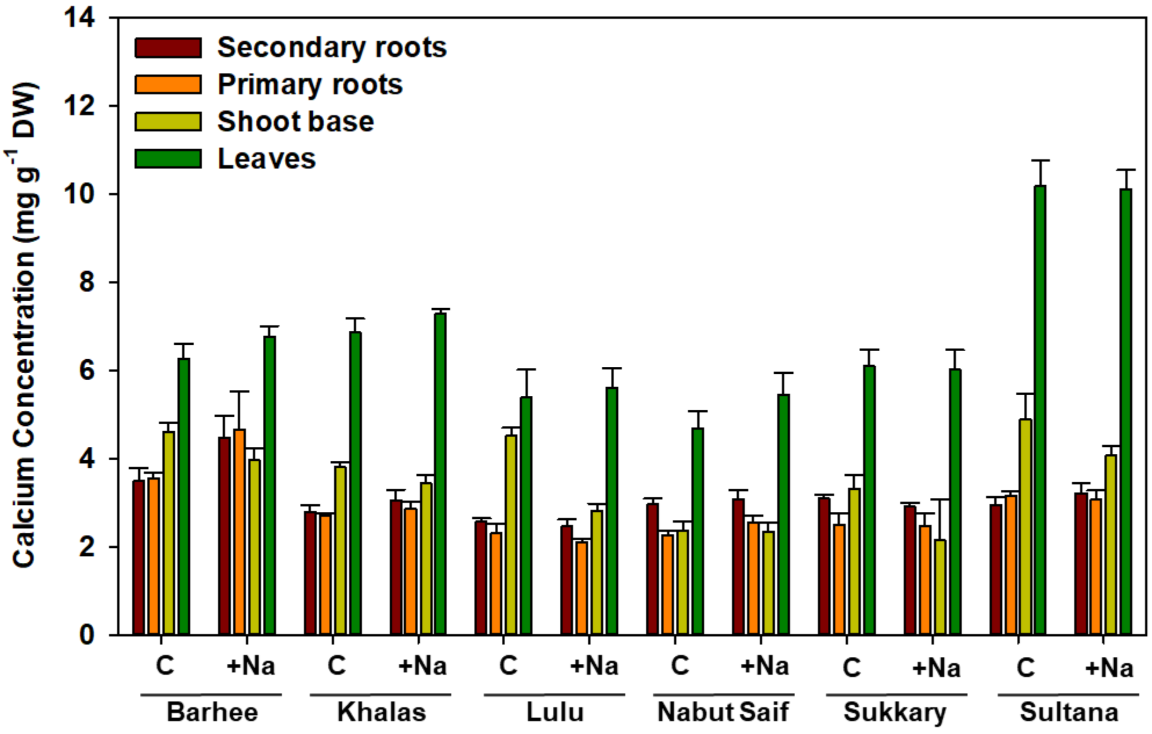


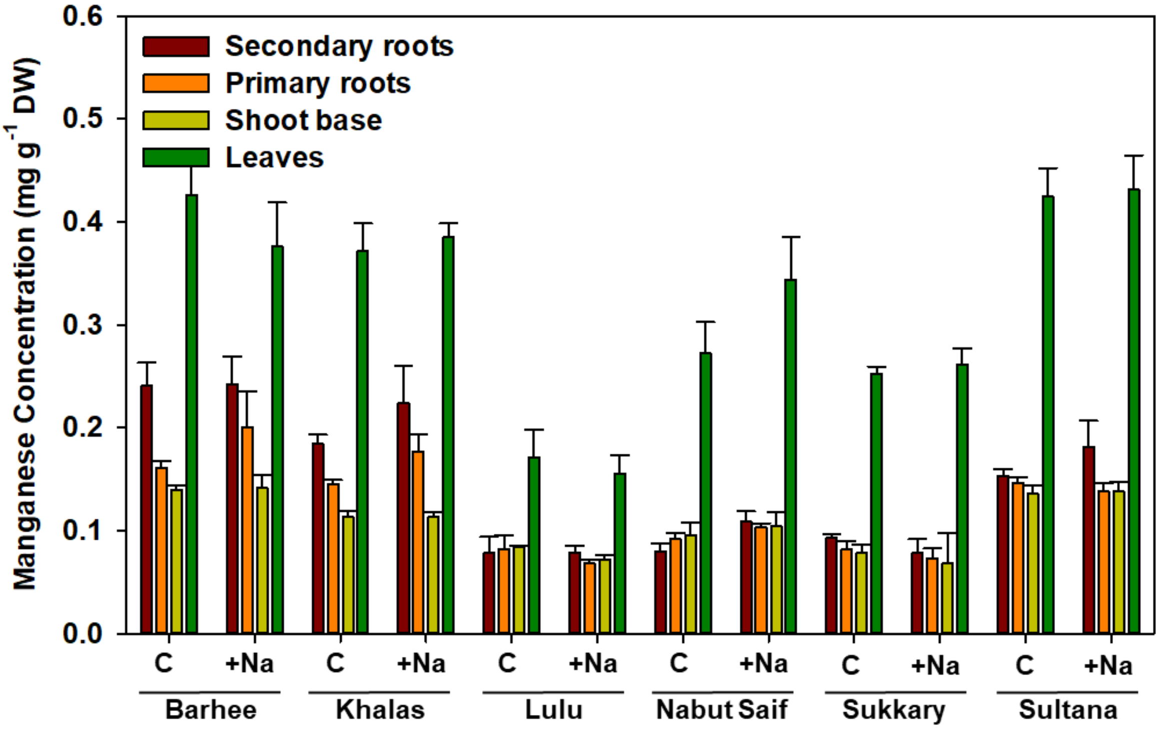


**Figure S4** Calcium and manganese concentration in different parts of six date palm (*Phoenix dactylifera*) varieties. Plants were irrigated with double-distilled water (Control, C) or once with 400 mL of 150 mM NaCl at the beginning of the experiment, then after two, four and six weeks with 400 mL of 300 mM NaCl (+Na); Experiment 1. Shown are averages ± standard errors. DW, dry weight.

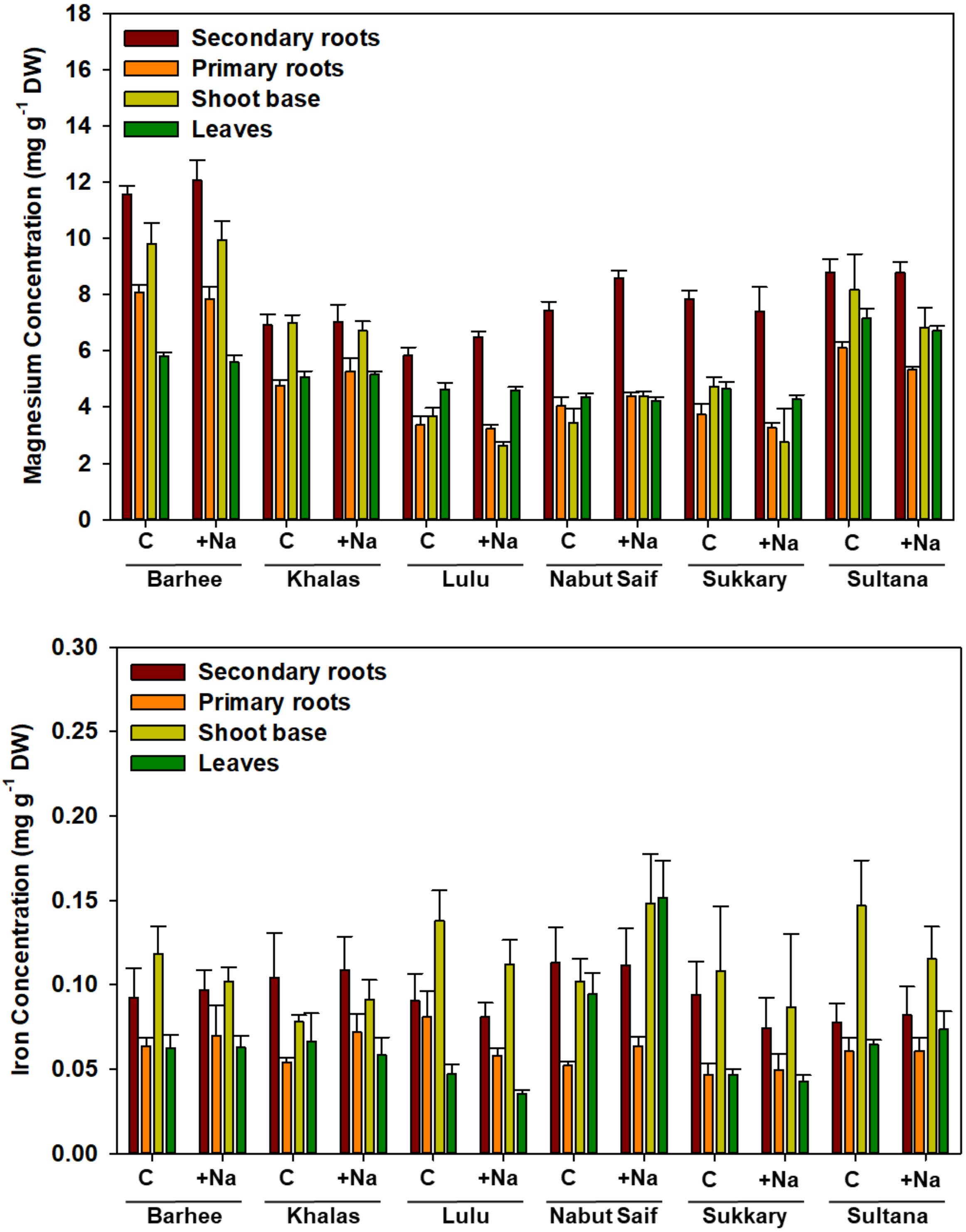


**Figure S5** Magnesium and iron concentration in different parts of six date palm (*Phoenix dactylifera*) varieties. Plants were irrigated with double-distilled water (Control, C) or once with 400 mL of 150 mM NaCl at the beginning of the experiment, then after two, four and six weeks with 400 mL of 300 mM NaCl (+Na); Experiment 1. Shown are averages ± standard errors. DW, dry weight.

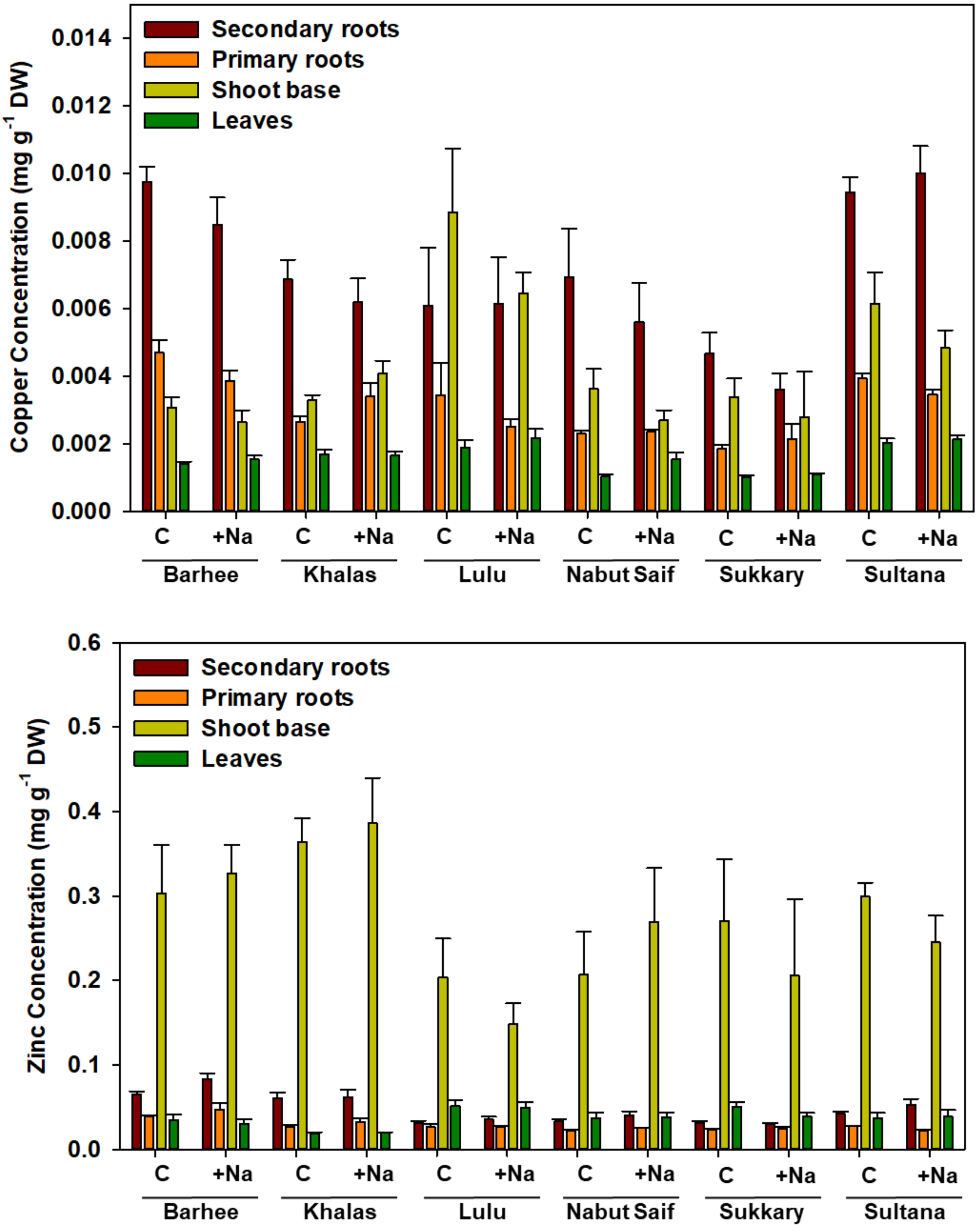


**Figure S6** Copper and zinc concentration in different parts of six date palm (*Phoenix dactylifera*) varieties. Plants were irrigated with double-distilled water (Control, C) or once with 400 mL of 150 mM NaCl at the beginning of the experiment, then after two, four and six weeks with 400 mL of 300 mM NaCl (+Na); Experiment 1. Shown are averages ± standard errors. DW, dry weight.

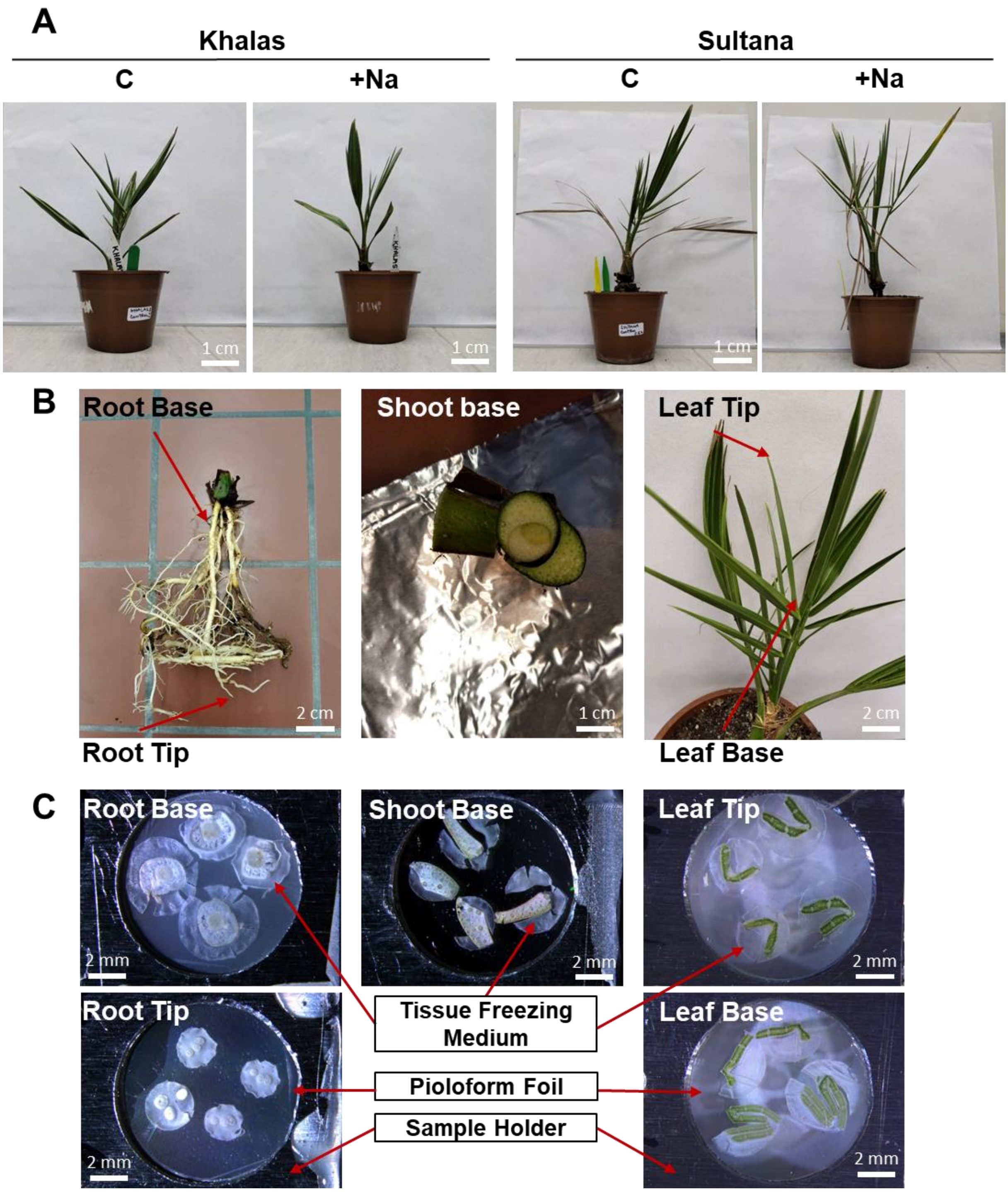


**Figure S7** Plants from Experiment 2 and samples for the localisation analysis. A: photographs of two date palm (*Phoenix dactylifera*) varieties, which were irrigated with double-distilled water (Control, C) or with 150 mM NaCl for two weeks and 300 mM NaCl for six weeks (+Na). B: Photographs of parts of plants sampled for cryo-fixation, cryo-sectioning and freeze-drying. C: representative freeze-dried cross-section of roots, shoot base and leaves sandwiched between two Pioloform foils stretched over aluminium sample holders.

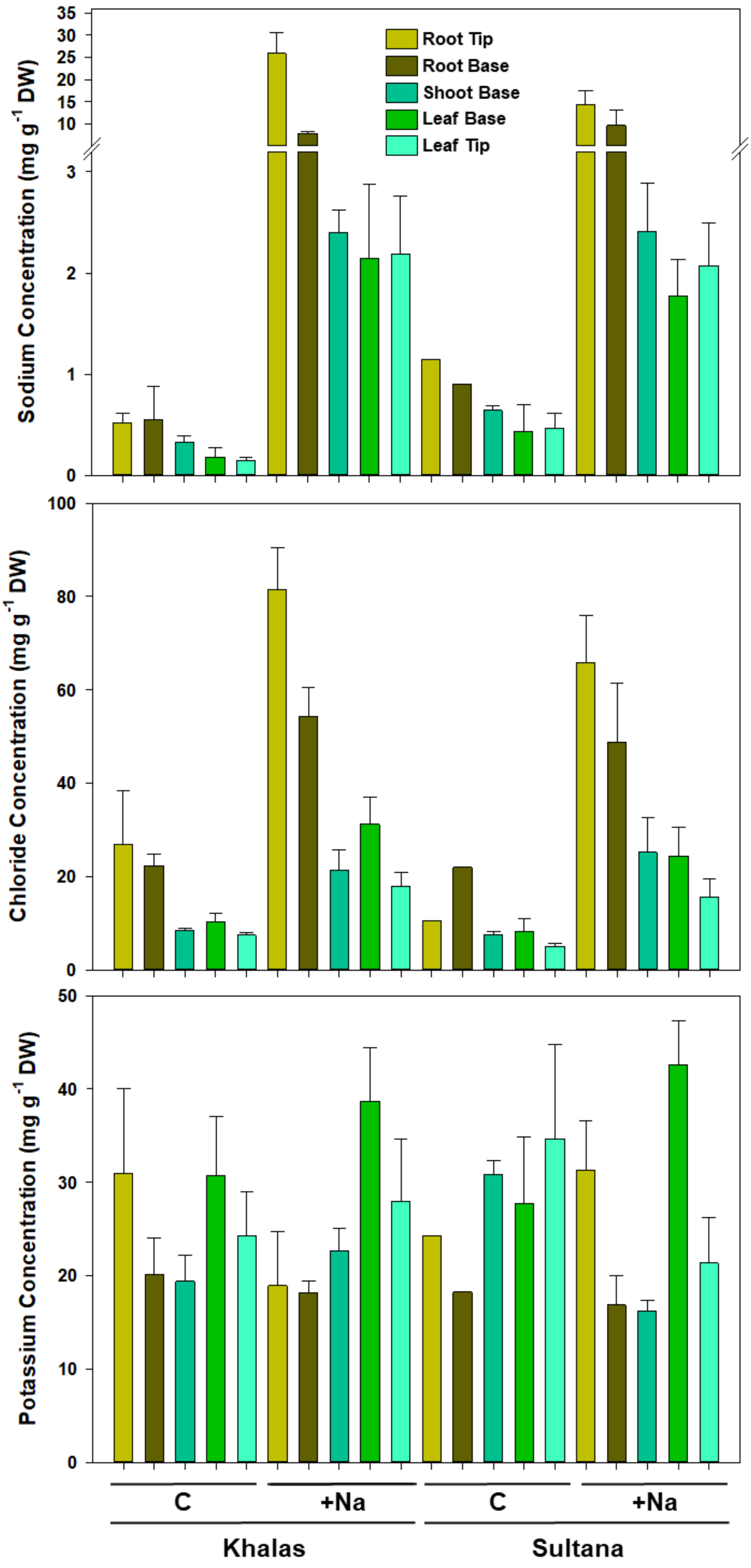


**Figure S8** Sodium, chlorine and potassium concentration in different parts of two date palm (*Phoenix dactylifera*) varieties. Plants were irrigated with double-distilled water (Control, C) or with 150 mM NaCl for two weeks and 300 mM NaCl for six weeks (+Na); Experiment 2. Shown are averages ± standard errors (n=1-3). DW, dry weight;

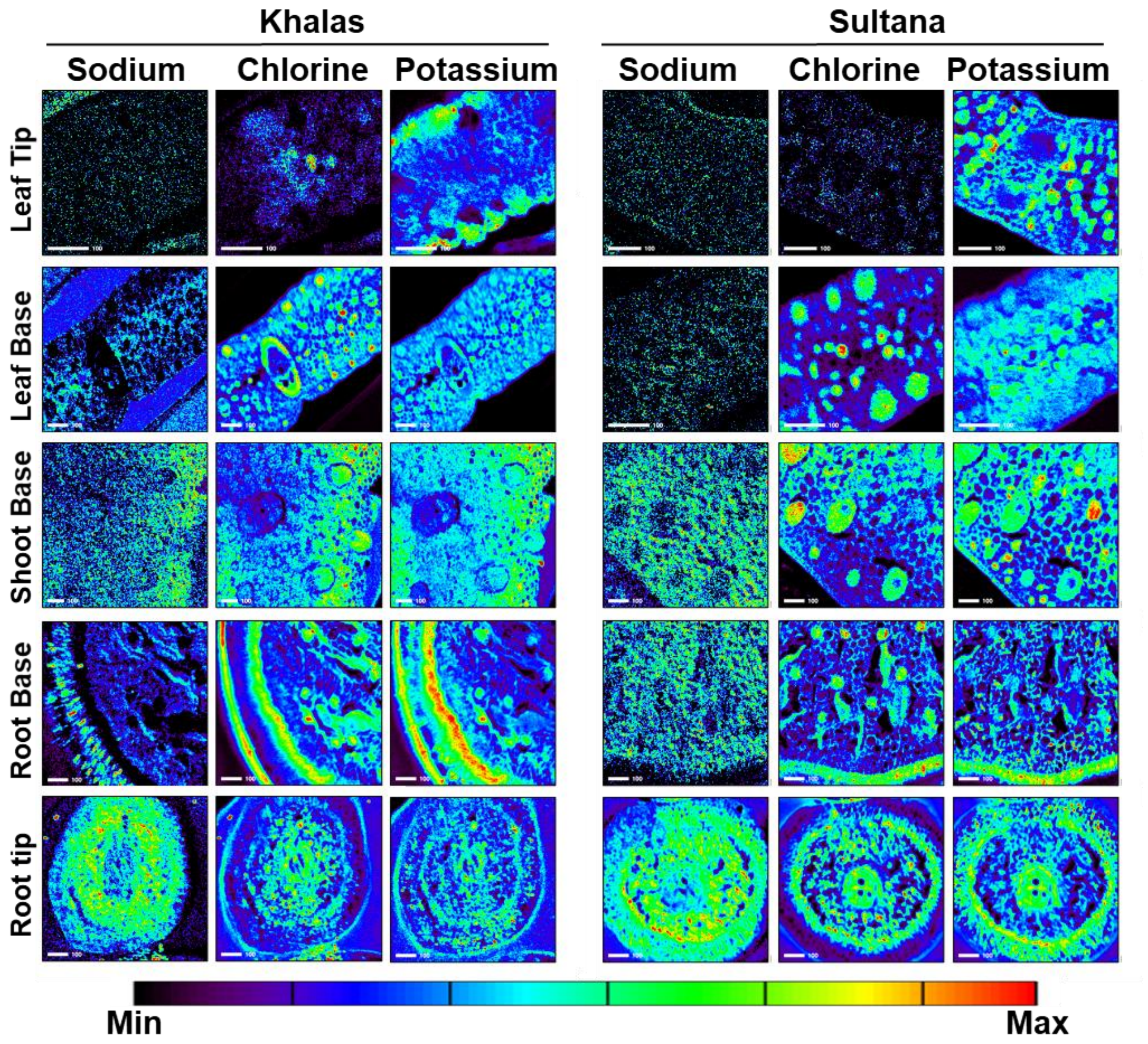


**Figure S9** Tissue-specific sodium, chlorine and potassium distribution in different parts of two date palm (*Phoenix dactylifera*) varieties. Plants were irrigated or with 150 mM NaCl for two weeks and 300 mM NaCl for six weeks; Experiment 2. Scale bars represent 100 µm.

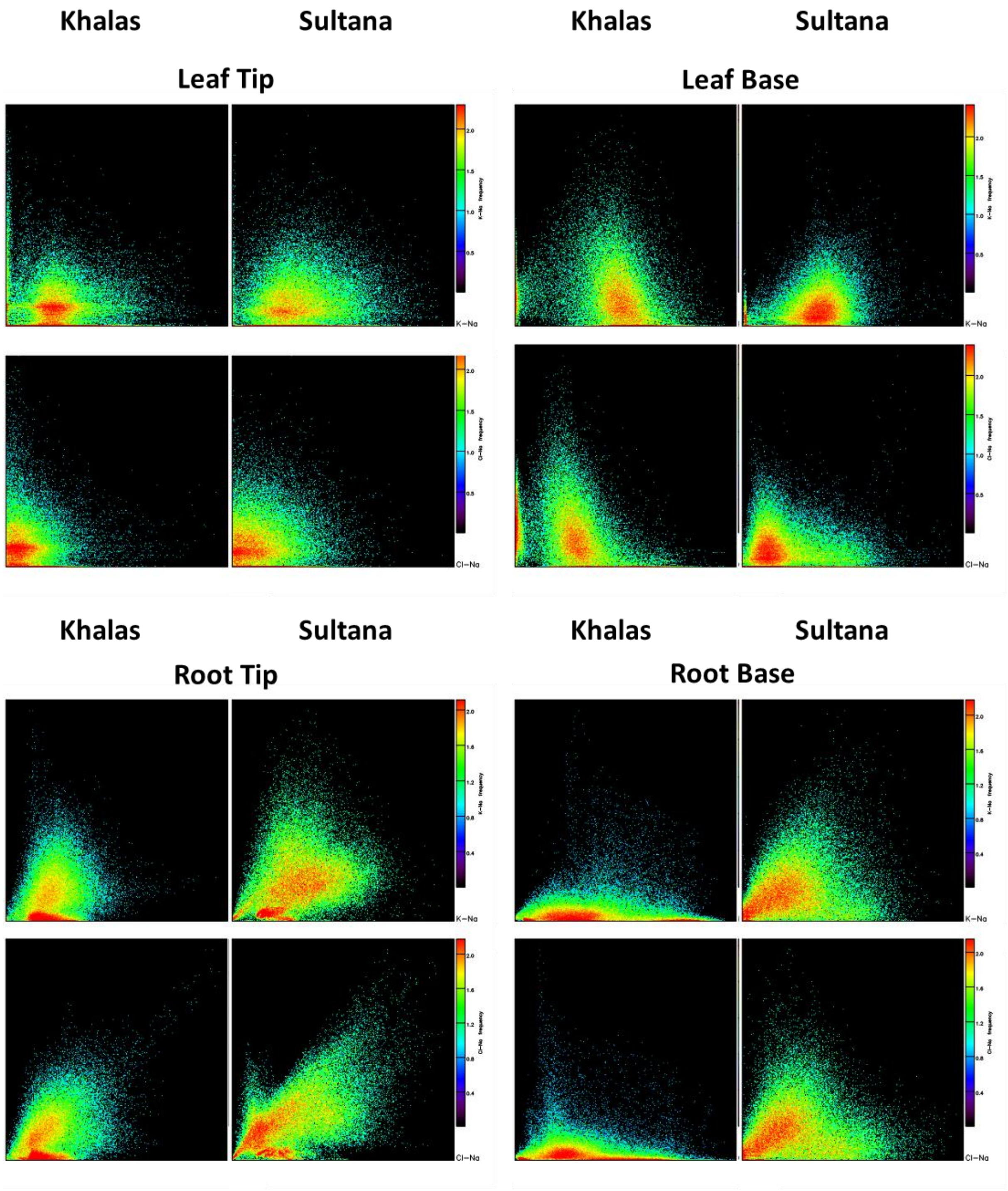


**Figure S10** Element association (correlation) of sodium (Na), potassium (K) and chlorine (Cl) concentration in their respective distribution maps (shown in Figure S9) in different parts of two date palm (*Phoenix dactylifera*) varieties. Plants were irrigated or with 150 mM NaCl for two weeks and 300 mM NaCl for six weeks; Experiment 2.

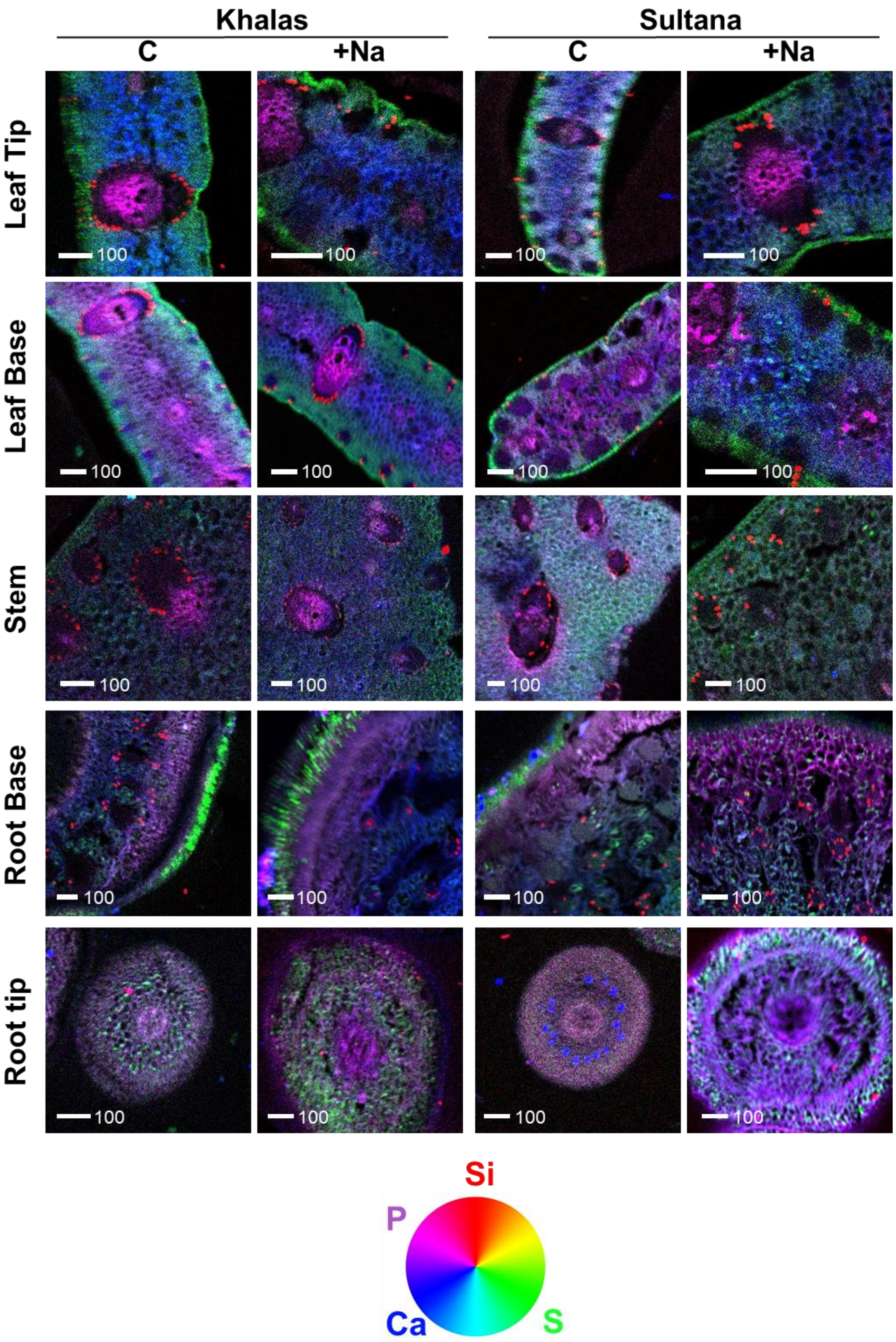


**Figure S11** Overlay of representative tissue-specific distribution maps of silicon (Si) in red, sulphur (S) in green, calcium (Ca) in blue and phosphorus (P) in magenta, in different parts of two date palm (*Phoenix dactylifera*) varieties. Plants were irrigated with double-distilled water (Control, C) or with 150 mM NaCl for two weeks and 300 mM NaCl for six weeks (+Na); Experiment 2. Scale bars are in µm.

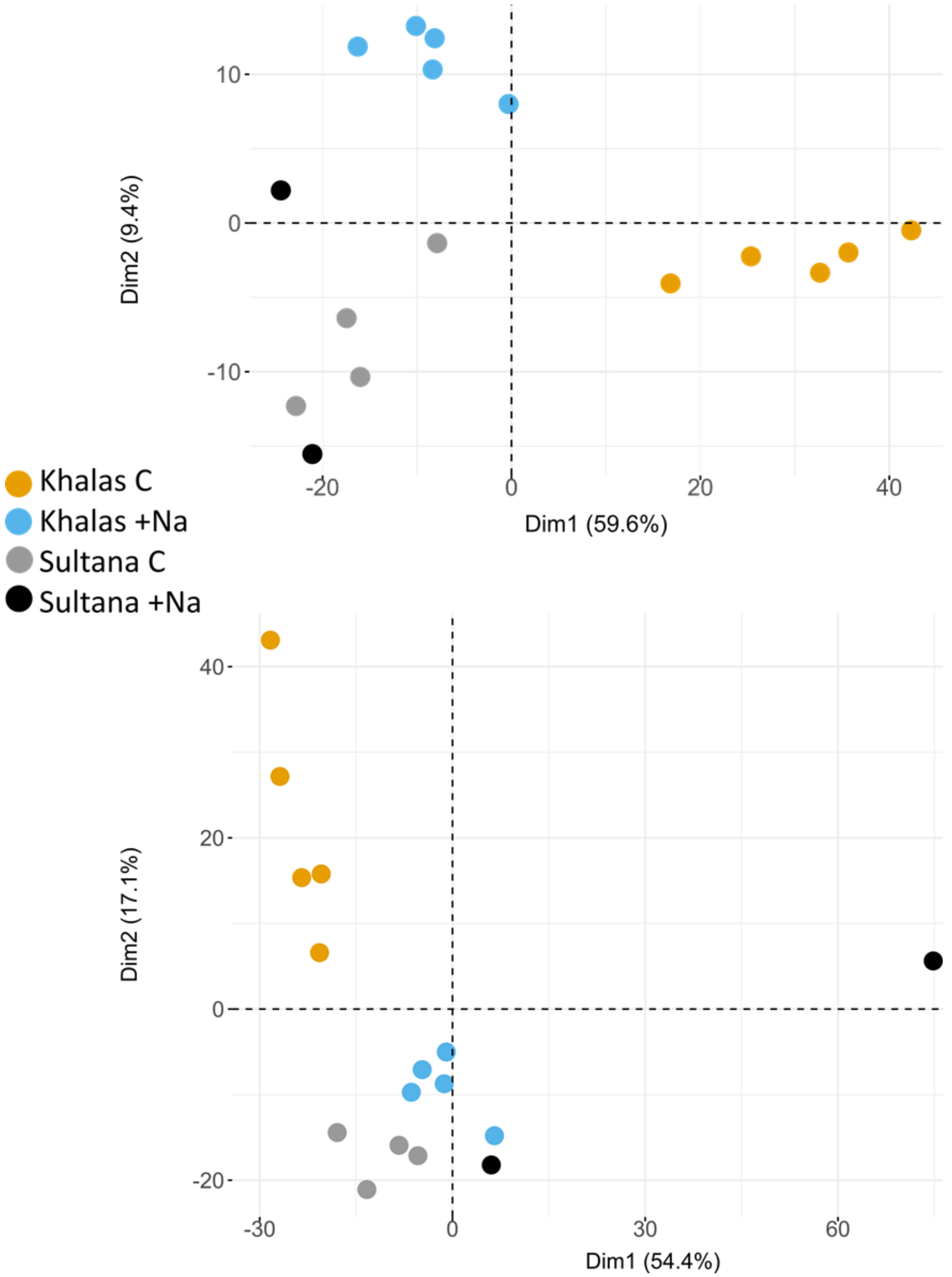


**Figure S12** Principal Component Analysis (PCA) of all differentially expressed genes in roots (top) and leaves (bottom) of two date palm (*Phoenix dactylifera*) varieties. Plants were irrigated with double-distilled water (Control, C) or with 150 mM NaCl for two weeks and 300 mM NaCl for six weeks (+Na). The PCA of 812 genes differentially expressed genes in roots in the +Na treatment showed clear clustering between Khalas and Sultana and between C and +Na treatment for Khalas but not for Sultana. The PCA of 1891 genes differentially expressed in leaves in the +Na treatment showed clear clustering between Khalas and Sultana and between C and +Na treatment for Khalas but not for Sultana.

**Table S1** Sodium (Na) and chloride (Cl) concentration (mg $g^{-1}$ dry weight) in roots (calculated as average concentration in secondary and primary roots) of two date palm (*Phoenix dactylifera*) varieties from Experiment 1. Plants were irrigated with double-distilled water (Control; C) or with NaCl for two weeks with 150 mM and for six weeks with 300 mM (+Na). n, number of repetitions; SE, standard error

| Variety | Treatment | | Na | Cl |
|---|---|---|---|---|
| **Barhee** | **C** | average | 1.058 | 10.5 |
| | | SE | 0.0375 | 0.695 |
| | | n | 6 | 6 |
| | **+Na** | average | 4.91 | 13.576 |
| | | SE | 0.381 | 0.411 |
| | | n | 6 | 6 |
| **Khalas** | **C** | average | 0.762 | 9.67 |
| | | SE | 0.0407 | 0.926 |
| | | n | 6 | 6 |
| | **+Na** | average | 3.128 | 12.928 |
| | | SE | 0.0558 | 0.748 |
| | | n | 6 | 6 |
| **Lulu** | **C** | average | 1.006 | 8.87 |
| | | SE | 0.0452 | 0.309 |
| | | n | 6 | 6 |
| | **+Na** | average | 5.635 | 11.92 |
| | | SE | 0.54 | 0.963 |
| | | n | 6 | 6 |
| **Nabut Saif** | **C** | average | 1.686 | 9.085 |
| | | SE | 0.0762 | 0.399 |
| | | n | 6 | 6 |
| | **+Na** | average | 7.319 | 12.022 |
| | | SE | 0.394 | 1.148 |
| | | n | 6 | 6 |
| **Sukkary** | **C** | average | 1.199 | 8.28 |
| | | SE | 0.0647 | 0.518 |
| | | n | 6 | 6 |
| | **+Na** | average | 5.349 | 11.434 |
| | | SE | 0.356 | 0.694 |
| | | n | 6 | 6 |
| **Sultana** | **C** | average | 0.593 | 9.67 |
| | | SE | 0.0201 | 0.488 |
| | | n | 6 | 6 |
| | **+Na** | average | 3.594 | 12.167 |
| | | SE | 0.153 | 0.937 |
| | | n | 6 | 6 |

**Table S2** Translocation factors (Leaves/ Roots concentration quotient) for essential elements in six date palm (*Phoenix dactylifera*) varieties from Experiment 1. Plants were irrigated with double-distilled water (Control) or once with 400 mL of 150 mM NaCl at the beginning of the experiment, then after two, four and six weeks with 400 mL of 300 mM NaCl (+Na). Mg, magnesium; P; phosphorus; S, sulphur; Ca, calcium; Mn, manganese; Fe, iron; Cu, copper; Zn, zinc

| Variety | Treatment | Mg | P | S | Ca | Mn | Fe | Cu | Zn |
|---|---|---|---|---|---|---|---|---|---|
| **Barhee** | **C** | 0.590 | 0.854 | 0.626 | 1.783 | 2.114 | 0.804 | 0.192 | 0.665 |
| | **+Na** | 0.563 | 0.929 | 0.613 | 1.481 | 1.698 | 0.754 | 0.248 | 0.470 |
| **Khalas** | **C** | 0.869 | 1.168 | 0.544 | 2.493 | 2.242 | 0.843 | 0.355 | 0.431 |
| | **+Na** | 0.839 | 1.203 | 0.368 | 2.465 | 1.920 | 0.644 | 0.344 | 0.411 |
| **Lulu** | **C** | 1.008 | 1.278 | 0.981 | 2.205 | 2.132 | 0.632 | 0.398 | 1.785 |
| | **+Na** | 0.942 | 1.203 | 0.937 | 2.452 | 2.137 | 0.531 | 0.502 | 1.595 |
| **Nabut Saif** | **C** | 0.757 | 1.690 | 0.554 | 1.788 | 3.168 | 1.141 | 0.223 | 1.341 |
| | **+Na** | 0.653 | 1.848 | 0.439 | 1.940 | 3.245 | 1.730 | 0.387 | 1.160 |
| **Sukkary** | **C** | 0.803 | 1.054 | 0.675 | 2.183 | 2.887 | 0.665 | 0.311 | 1.868 |
| | **+Na** | 0.803 | 1.042 | 0.580 | 2.242 | 3.459 | 0.693 | 0.377 | 1.430 |
| **Sultana** | **C** | 0.962 | 1.643 | 1.117 | 3.330 | 2.836 | 0.934 | 0.304 | 1.079 |
| | **+Na** | 0.950 | 1.543 | 1.055 | 3.216 | 2.694 | 1.029 | 0.316 | 1.031 |
| **Overall average** | **C** | 0.832 | 1.281 | 0.750 | 2.297 | 2.563 | 0.836 | 0.297 | 1.195 |
| | **+Na** | 0.792 | 1.295 | 0.665 | 2.299 | 2.525 | 0.897 | 0.363 | 1.016 |

**Table S3** Element concentration (mg $g^{-1}$ dry weight) in different parts of two date palm (*Phoenix dactylifera*) varieties and translocation factors (Leaves/Roots concentration quotient) from Experiment 2. Plants were irrigated with double-distilled water (Control) or with NaCl for two weeks with 150 mM and for six weeks with 300 mM (+Na). Mg, magnesium; P; phosphorus; S, sulphur; Ca, calcium; Mn, manganese; Fe, iron; Cu, copper; Ni, nickel; Zn, zinc; n, number of repetitions; SE, standard error

| | **var. Khalas** | | **Mg** | **P** | **S** | **Ca** | **Mn** | **Fe** | **Cu** | **Zn** |
|---|---|---|---|---|---|---|---|---|---|---|
| **Control** | **Roots** | Mean | 2.495 | 2.701 | 2.988 | 4.583 | 0.0215 | 0.0568 | 0.00485 | 0.026 |
| | | SE | 0.257 | 0.0892 | 0.178 | 0.816 | 0.00141 | 0.00341 | 0.000333 | 0.00516 |
| | | n | 2 | 2 | 2 | 2 | 2 | 2 | 2 | 2 |
| | **Root Tips** | Mean | 4.435 | 2.851 | 5.397 | 4.176 | 0.0194 | 0.156 | -- | 0.0344 |
| | | SE | -- | -- | -- | -- | -- | -- | -- | -- |
| | | n | 1 | 1 | 1 | 1 | 1 | 1 | 0 | 1 |
| | **Root Base** | Mean | 2.457 | 3.903 | 3.869 | 4.062 | 0.0162 | 0.0795 | 0.00564 | 0.0235 |
| | | SE | 0.556 | 1.32 | 0.457 | 0.519 | 0.00409 | 0.0324 | 0.00276 | 0.00911 |
| | | n | 2 | 2 | 2 | 2 | 2 | 2 | 2 | 2 |
| | **Shoot Base** | Mean | 1.481 | 2.746 | 2.474 | 2.046 | 0.0264 | 0.0302 | 0.00298 | 0.0901 |
| | | SE | 0.291 | 0.392 | 0.355 | 0.306 | 0.00609 | 0.00503 | 0.000208 | 0.054 |
| | | n | 4 | 4 | 4 | 4 | 4 | 4 | 4 | 4 |
| | **Leaves** | Mean | 1.164 | 2.366 | 2.221 | 3.023 | 0.191 | 0.0574 | 0.0031 | 0.0258 |
| | | SE | 0.0997 | 0.0684 | 0.171 | 0.253 | 0.0609 | 0.0147 | 0.000249 | 0.00682 |
| | | n | 2 | 2 | 2 | 2 | 2 | 2 | 2 | 2 |
| | **Leaf Base** | Mean | 1.161 | 3.114 | 2.065 | 2.048 | 0.0579 | 0.0363 | 0.00364 | 0.0231 |
| | | SE | 0.252 | 0.316 | 0.0185 | 0.0143 | 0.0112 | 0.00237 | 0.000963 | 0.00081 |
| | | n | 2 | 2 | 2 | 2 | 2 | 2 | 2 | 2 |
| | **Leaf Tip** | Mean | 2.526 | 1.341 | 2.721 | 7.749 | 0.274 | 0.118 | 0.01 | 0.0238 |
| | | SE | 0.362 | 0.214 | 0.469 | 0.0749 | 0 | 0.0616 | 0.00192 | 0.00308 |
| | | n | 2 | 2 | 2 | 2 | 1 | 2 | 2 | 2 |
| **+Na** | **Roots** | Mean | 2.89 | 3.673 | 3.785 | 9.247 | 0.0757 | 0.111 | 0.00862 | 0.047 |
| | | SE | 0.0523 | 0.703 | 0.132 | 3.911 | 0.0138 | 0.0118 | 0.00124 | 0.00824 |
| | | n | 2 | 2 | 2 | 2 | 2 | 2 | 2 | 2 |
| | **Root Tips** | Mean | 7.227 | 5.352 | 5.899 | 4.051 | 0.0578 | 0.0562 | 0.00375 | 0.0581 |
| | | SE | 0 | 0 | 0 | 0 | 0 | 0 | 0 | 0 |
| | | n | 1 | 1 | 1 | 1 | 1 | 1 | 1 | 1 |
| | **Root Base** | Mean | 1.821 | 3.23 | 3.326 | 3.118 | 0.0233 | 0.0253 | 0.00235 | 0.0122 |
| | | SE | 0.285 | 0.28 | 0.258 | 0.0366 | 0.00533 | 0.00337 | 0.00052 | 0.00121 |
| | | n | 2 | 2 | 2 | 2 | 2 | 2 | 2 | 2 |
| | **Shoot Base** | Mean | 1.465 | 2.626 | 2.779 | 2.437 | 0.0318 | 0.025 | 0.00212 | 0.0352 |
| | | SE | 0.206 | 0.209 | 0.127 | 0.188 | 0.00287 | 0.00228 | 0.000204 | 0.0156 |
| | | n | 4 | 4 | 4 | 4 | 4 | 4 | 4 | 4 |
| | **Leaves** | Mean | 1.237 | 3.038 | 2.652 | 4.315 | 0.188 | 0.0451 | 0.00221 | 0.0252 |
| | | SE | 0.00726 | 0.314 | 0.213 | 0.725 | 0.0821 | 0.00784 | 0.000448 | 0.00391 |
| | | n | 2 | 2 | 2 | 2 | 2 | 2 | 2 | 2 |
| | **Leaf Base** | Mean | 1.01 | 3.379 | 2.215 | 2.695 | 0.0475 | 0.0404 | 0.00346 | 0.0291 |
| | | SE | 0.263 | 0.437 | 0.106 | 0.713 | 0.00824 | 0.000558 | 0.000541 | 0.000895 |
| | | n | 2 | 2 | 2 | 2 | 2 | 2 | 2 | 2 |
| | **Leaf Tip** | Mean | 2.16 | 1.738 | 2.982 | 7.324 | 0.237 | 0.0393 | 0.0024 | 0.0191 |
| | | SE | 0.559 | 0.172 | 0.379 | 0.0275 | 0.0624 | 0.00239 | 0.000509 | 0.00264 |
| | | n | 2 | 2 | 2 | 2 | 2 | 2 | 2 | 2 |
| | **Leaves/ Roots** | **C** | 0.467 | 0.876 | 0.743 | 0.660 | 8.88 | 1.011 | 0.639 | 0.992 |
| | | **+Na** | 0.428 | 0.827 | 0.701 | 0.467 | 2.483 | 0.406 | 0.256 | 0.536 |

**Table S3** Continued…

| var. Sultana | | | Mg | P | S | Ca | Mn | Fe | Cu | Zn |
|---|---|---|---|---|---|---|---|---|---|---|
| Control | Roots | Mean | 5.3 | 3.002 | 4.279 | 14.849 | 0.0817 | 0.111 | 0.019 | 0.0529 |
| | | SE | 0 | 0 | 0 | 0 | 0 | 0 | 0 | 0 |
| | | n | 1 | 1 | 1 | 1 | 1 | 1 | 1 | 1 |
| | Root Tips | Mean | -- | -- | -- | -- | -- | -- | -- | -- |
| | | SE | -- | -- | -- | -- | -- | -- | -- | -- |
| | | n | 0 | 0 | 0 | 0 | 0 | 0 | 0 | 0 |
| | Root Base | Mean | 2.39 | 2.632 | 4.536 | 5.892 | 0.0296 | 0.0668 | 0.0108 | 0.0344 |
| | | SE | 0 | 0 | 0 | 0 | 0 | 0 | 0 | 0 |
| | | n | 1 | 1 | 1 | 1 | 1 | 1 | 1 | 1 |
| | Shoot Base | Mean | 2.017 | 3.721 | 4.836 | 4.143 | 0.074 | 0.0702 | 0.00526 | 0.037 |
| | | SE | 0.698 | 0.723 | 1.086 | 2.153 | 0.0243 | 0.0236 | 0.00154 | 0.00715 |
| | | n | 4 | 4 | 4 | 4 | 4 | 4 | 4 | 4 |
| | Leaves | Mean | 1.074 | 2.571 | 3.321 | 2.665 | 0.109 | 0.0296 | 0.00237 | 0.0186 |
| | | SE | 0.346 | 0.122 | 0.469 | 0.933 | 0.042 | 0.00269 | 0.000413 | 0.00193 |
| | | n | 2 | 2 | 2 | 2 | 2 | 2 | 2 | 2 |
| | Leaf Base | Mean | 2.362 | 3.265 | 4.12 | 4.7 | 0.0664 | 0.0368 | 0.0046 | 0.12 |
| | | SE | 1.469 | 0.0275 | 1.757 | 3.107 | 4.51E-05 | 0.00179 | 0.00168 | 0.0969 |
| | | n | 2 | 2 | 2 | 2 | 2 | 2 | 2 | 2 |
| | Leaf Tip | Mean | 1.707 | 1.288 | 3.435 | 5.1 | 0.219 | 0.0427 | 0.00238 | 0.0191 |
| | | SE | 0.129 | 0.252 | 0.028 | 0.29 | 0.117 | 0.0128 | 0.000447 | 0.000236 |
| | | n | 2 | 2 | 2 | 2 | 2 | 2 | 2 | 2 |
| +Na | Roots | Mean | 2.486 | 2.655 | 5.902 | 5.668 | 0.0713 | 0.122 | 0.0178 | 0.0438 |
| | | SE | 0.0444 | 0.203 | 0.551 | 0.144 | 0.0107 | 0.0483 | 0.00425 | 0.00586 |
| | | n | 3 | 3 | 3 | 3 | 2 | 2 | 2 | 3 |
| | Root Tips | Mean | 6.791 | 4.631 | 10.659 | 4.031 | 0.0905 | 0.0655 | 0.00645 | 0.0593 |
| | | SE | 1.741 | 1.985 | 2.077 | 0.792 | 0.0399 | 0.0109 | 0.00158 | 0.0307 |
| | | n | 3 | 3 | 3 | 3 | 3 | 3 | 3 | 3 |
| | Root Base | Mean | 2.308 | 2.966 | 5.477 | 3.426 | 0.0741 | 0.0453 | 0.00478 | 0.0277 |
| | | SE | 0.373 | 0.437 | 0.283 | 0.59 | 0.0344 | 0.00837 | 0.00197 | 0.00918 |
| | | n | 3 | 3 | 3 | 3 | 3 | 3 | 3 | 3 |
| | Shoot Base | Mean | 1.995 | 2.861 | 3.612 | 2.397 | 0.0742 | 0.0358 | 0.00537 | 0.0427 |
| | | SE | 0.843 | 0.453 | 0.969 | 0.74 | 0.0319 | 0.00909 | 0.0026 | 0.0204 |
| | | n | 6 | 6 | 6 | 5 | 6 | 5 | 6 | 6 |
| | Leaves | Mean | 0.868 | 3.961 | 2.984 | 3.787 | 0.132 | 0.0483 | 0.00243 | 0.0267 |
| | | SE | 0.067 | 0.506 | 0.522 | 0.31 | 0.0133 | 0.00595 | 0.000492 | 0.00394 |
| | | n | 3 | 3 | 3 | 3 | 3 | 3 | 3 | 3 |
| | Leaf Base | Mean | 0.789 | 4.845 | 2.477 | 2.982 | 0.0906 | 0.0512 | 0.00222 | 0.0248 |
| | | SE | 0.0571 | 1.07 | 0.264 | 0.906 | 0.0286 | 0.00289 | 0.000318 | 0.00585 |
| | | n | 3 | 3 | 3 | 3 | 3 | 3 | 3 | 3 |
| | Leaf Tip | Mean | 1.591 | 2.892 | 3.103 | 7.419 | 0.359 | 0.0546 | 0.00257 | 0.0244 |
| | | SE | 0.127 | 0.759 | 0.178 | 0.861 | 0.12 | 0.0115 | 0.000673 | 0.00336 |
| | | n | 3 | 3 | 3 | 3 | 2 | 3 | 3 | 3 |
| | Leaves/ Roots | C | 0.203 | 0.856 | 0.776 | 0.179 | 1.33 | 0.267 | 0.125 | 0.352 |
| | | +Na | 0.349 | 1.492 | 0.506 | 0.668 | 1.851 | 0.396 | 0.137 | 0.610 |

**Table Captions for Tables S4-12 available at https://doi.org/10.5281/zenodo.14283064**

**Table S4** Differentially expressed genes (DEGs) in roots of a date palm (*Phoenix dactylifera*) variety Khalas (K). Plants were watered either with double-distilled water (C) or with NaCl (Na) for two weeks with 150 mM and for six weeks with 300 mM; Experiment 3.

**Table S5** Differentially expressed genes (DEGs) in shoots of a date palm (*Phoenix dactylifera*) variety Khalas (K). Plants were watered either with double-distilled water (C) or with NaCl (Na) for two weeks with 150 mM and for six weeks with 300 mM; Experiment 3.

**Table S6** Differentially expressed genes (DEGs) in roots of a date palm (*Phoenix dactylifera*) variety Sultana (S). Plants were watered either with double-distilled water (C) or with NaCl (Na) for two weeks with 150 mM and for six weeks with 300 mM; Experiment 3.

**Table S7** Differentially expressed genes (DEGs) in shoots of a date palm (*Phoenix dactylifera*) variety Sultana (S). Plants were watered either with double-distilled water (C) or with NaCl (Na) for two weeks with 150 mM and for six weeks with 300 mM; Experiment 3.

**Table S8** Differentially expressed genes (DEGs) in roots of two date palm (*Phoenix dactylifera*) varieties Khalas (K) and Sultana (S). Plants were watered with double-distilled water (C) for eight weeks; Experiment 3.

**Table S9** Differentially expressed genes (DEGs) in shoots of two date palm (*Phoenix dactylifera*) varieties Khalas (K) and Sultana (S). Plants were watered with double-distilled water (C) for eight weeks; Experiment 3.

**Table S10** Differentially expressed genes (DEGs) in roots of two date palm (*Phoenix dactylifera*) varieties Khalas (K) and Sultana (S). Plants were watered with NaCl (Na) for two weeks with 150 mM and for six weeks with 300 mM; Experiment 3.

**Table S11** Differentially expressed genes (DEGs) in shoots of two date palm (*Phoenix dactylifera*) varieties Khalas (K) and Sultana (S). Plants were watered with NaCl (Na) for two weeks with 150 mM and for six weeks with 300 mM; Experiment 3.

**Table S12** Gene ontology enrichment analysis. P-value cut off at 0.05. Two date palm (*Phoenix dactylifera*) varieties Khalas and Sultana. Plants were watered with NaCl (Na) for two weeks with 150 mM and for six weeks with 300 mM; Experiment 3.
